# NREM sleep stages specifically alter dynamical integration of large-scale brain networks

**DOI:** 10.1101/2020.07.08.193508

**Authors:** Anjali Tarun, Danyal Wainstein-Andriano, Virginie Sterpenich, Laurence Bayer, Lampros Perogamvros, Mark Solms, Nikolai Axmacher, Sophie Schwartz, Dimitri Van De Ville

## Abstract

Functional dissociations in the brain observed during non-rapid eye movement (NREM) sleep have been mainly associated with reduced information integration and impaired consciousness that accompany increasing sleep depth. Most of the studies that evaluated this relation have mostly been focused on *spatial* alterations of brain networks across various vigilance states. Here, we explored the *dynamical* properties of large-scale functional brain networks derived from *transients* or moments of activity changes in fMRI using two complementary studies of simultaneous EEG-fMRI recordings of sleep. Our results revealed distinct alterations in the individual network’s dynamical characteristics across wakefulness and NREM sleep. Positive activations of visual-sensory areas simultaneously deactivate focal subcortical regions known to be involved with arousal regulation. The cerebellum is also found to dissociate into posterior and anterior regions, with the former being predominantly present during wakefulness than in the deep sleep. Most surprisingly, we found that global network activity and pair-wise network interactions increased significantly in NREM stage 2 before an abrupt loss of communication is observed in NREM stage 3. Thus, by providing a spatiotemporal and more accurate network-level representation of brain organization, we were able to capture new features of information integration of consciousness during sleep, and provide concrete evidence for the presence of unstable yet distributed global synchronization in NREM stage 2.

## 1. Introduction

Spontaneous brain activity, as assessed by resting-state functional Magnetic Resonance Imaging (fMRI), has provided key insights into the functional architecture of the brain. Resting-state networks (RSNs) identify sets of brain regions that exhibit synchronized fluctuations of activity over the whole duration of a resting-state session (typically 10-20 min of continuous scanning). The main hypothesis underlying most functional connectivity (FC) studies is that different RSNs reflect distinct ongoing cognitive/affective processes/states. For instance, the default mode network (DMN) typically shows reduced activity when subjects perform an externally oriented task (Greicius et al., 2003) and, contrastingly, the DMN becomes more engaged when self-referential processes or internal mentation predominate (Andrews-Hanna, 2012). According to this interpretation of RSNs, we expect that if conscious awareness dissipates as the brain transitions from wakefulness to deep sleep, we should observe a parallel and net decrease in activity and/or FC across regions of the brain involved in higher-order cognitive operations, such as reasoning, monitoring, or metacognition. Yet, some early evidence also suggested that several RSNs, including those encompassing association networks, persisted or even increased their connectivity during the descent from wakefulness to light sleep (Larson-Prior et al., 2009) or during anesthesia and coma, when conscious awareness is presumed to be completely abolished (Boly et al., 2008). Therefore, RSNs may reflect intrinsic dynamical properties of the brain’s functional organization that are maintained across distinct levels of consciousness.

From a behavioral point of view, the brain in sleep undergoes marked and well-characterized physiological changes. Based on polysomnography (PSG), which is a combined use of EEG, electro-oculography, and electromyography, natural sleep can be broadly divided into rapid eye movement (REM) and non-rapid eye movement (NREM) periods. The latter is further subdivided into different sleep stages characterized by relaxed wakefulness (N1) to light sleep (N2), up to slow-wave sleep (SWS) or deep sleep (N3). It is therefore not surprising that, although RSNs may be detected across different sleep stages, their connectivity patterns (Horovitz et al., 2009; Sämann et al., 2011) and FC strengths undergo significant modifications (Tagliazucchi et al., 2013). For instance, upon reaching N3, the DMN has been found to dissociate into subcomponents, with a decrease in the connectivity between the medial prefrontal cortex (MPFC) and the posterior cingulate cortex (PCC) (Horovitz et al., 2009). Furthermore, several studies looked at changes in FC for regions associated with the reticular activation system (RAS) that regulate the physiological state of arousal during sleep (e.g. thalamus, hypothalamus) and reported decreased connectivity between these regions and the rest of the cortex during light and deep sleep (Hale et al., 2016; Picchioni et al., 2014; Tagliazucchi and Laufs, 2014). These changes in brain network integrity, particularly its marked reduction from wakefulness to deep sleep, have been associated to diminished level of information integration (Tononi, 2004). That is, when the brain switches to more local cortical processing, this would lead to a global loss of information integration and a concomitant reduction in consciousness (Tononi and Koch, 2015).

Methodological tools allowing the detection of prevalent brain spatial patterns, such as FC analyses, are particularly useful when assessing imaging data collected outside of a predefined experimental paradigm (or in the absence of any temporally-defined independent variable), namely when regression techniques cannot be applied. This is typically the case for continuous data collected during varying levels of arousal or states of consciousness (e.g. resting-state, sleep, anesthesia, and drugs). The simplest approach to investigating changes in FC is to use a sliding-window technique, where time-courses from sets of brain regions (e.g. from atlas-based parcellation) are segmented into successive temporal windows so that various assessments of FC (e.g. bivariate Pearson correlations) can be applied to obtain time-evolving connectivity matrices. Another approach is to derive analogous information on resting-state FC based on time-points where the regional BOLD signal exceeds a particular threshold (Tagliazucchi et al., 2012). Temporal clustering can also be applied to activity patterns occurring at these active fMRI time-points to obtain patterns of co-activity among regions, also known as co-activation patterns (CAPs) (Liu and Duyn, 2013).

Furthermore, to account for the fundamental dynamic nature of the changes in neural FC, the latest developments on non-stationary FC approaches have started to successfully incorporate methods of temporal modeling. Using a dynamic Bayesian approach (i.e., Hidden Markov Model; HMM), Stevner and colleagues (2019) could recently demonstrate that some specific whole-brain functional connections are associated to each of the different stages of NREM. They found that networks with high specificity to occur in stages N2 and N3 generally expressed longer mean lifetimes, with each HMM state lasting from a few seconds to tens of seconds. They also examined the transition probabilities between the networks by extracting modules of HMM states that transitioned more often between each other than to other states (Vidaurre et al., 2017), allowing them to identify key trajectories of network activity from wake to NREM sleep.

Despite these major methodological advancements, it remains unsettling that coordinated network activity is assumed to be strictly temporally segregated, and that whole-brain states are not overlapping in time (i.e., only one RSN can occur per time instance). To overcome this limitation, we adopted a new framework called innovation-driven co-activation patterns (iCAPS) which capture *transient* brain activity, *i.e.,* physiologically significant moments of regional activation and deactivation (Karahanoğlu and Van De Ville, 2015). This allows for the recovery of RSNs that are both spatially and temporally overlapping, providing a more plausible and thus putatively more accurate description of functional brain organization.

Furthermore, unlike standard CAPs and other data-driven approaches (e.g., independent component analysis; Smith et al. 2012), the iCAP approach explicitly accounts for temporal blurring by the hemodynamic response function (HRF) by incorporating a deconvolution step in the preprocessing of the fMRI signal. Using these advantages, we demonstrate that RSNs such as control, salience, visual and sensory networks persist until deep sleep, and showed that some networks naturally dissociate into subcomponents (*e.g.,* DMN, cerebellum). Moreover, we expect to observe many networks that overlap in time, and that these numbers are altered across the different levels of sleep. To test these hypotheses, temporal measures and pair-wise network interactions such as overall accumulated durations, mean lifetimes, couplings, and temporal overlaps were computed. We put the focal point into detecting the *temporal* alterations of large-scale brain activity across the distinct vigilance states. Finally, we show how specific changes in the dynamical interactions of large-scale brain networks can capture new features of information integration theory of consciousness.

## 2. Results

### 2.1 Data distribution across Study 1 and Study 2

Fig. 1 illustrates the two experimental paradigms on simultaneously acquired EEG and fMRI recordings of sleep. We used a total of 21 subjects in Study 1 (13 reached N3), and a total of 7 subjects in the Study 2 (all of them reached N1 sleep). The distribution of accumulated data from each sleep stage for each dataset is also shown in Fig. 1, with Study 1 generating a substantial amount of N3 sleep (more than 8 hours in total), while Study 2 was predominated by wake and N1 sleep (about 8h and 4h of data, respectively).

**Fig. 1.**
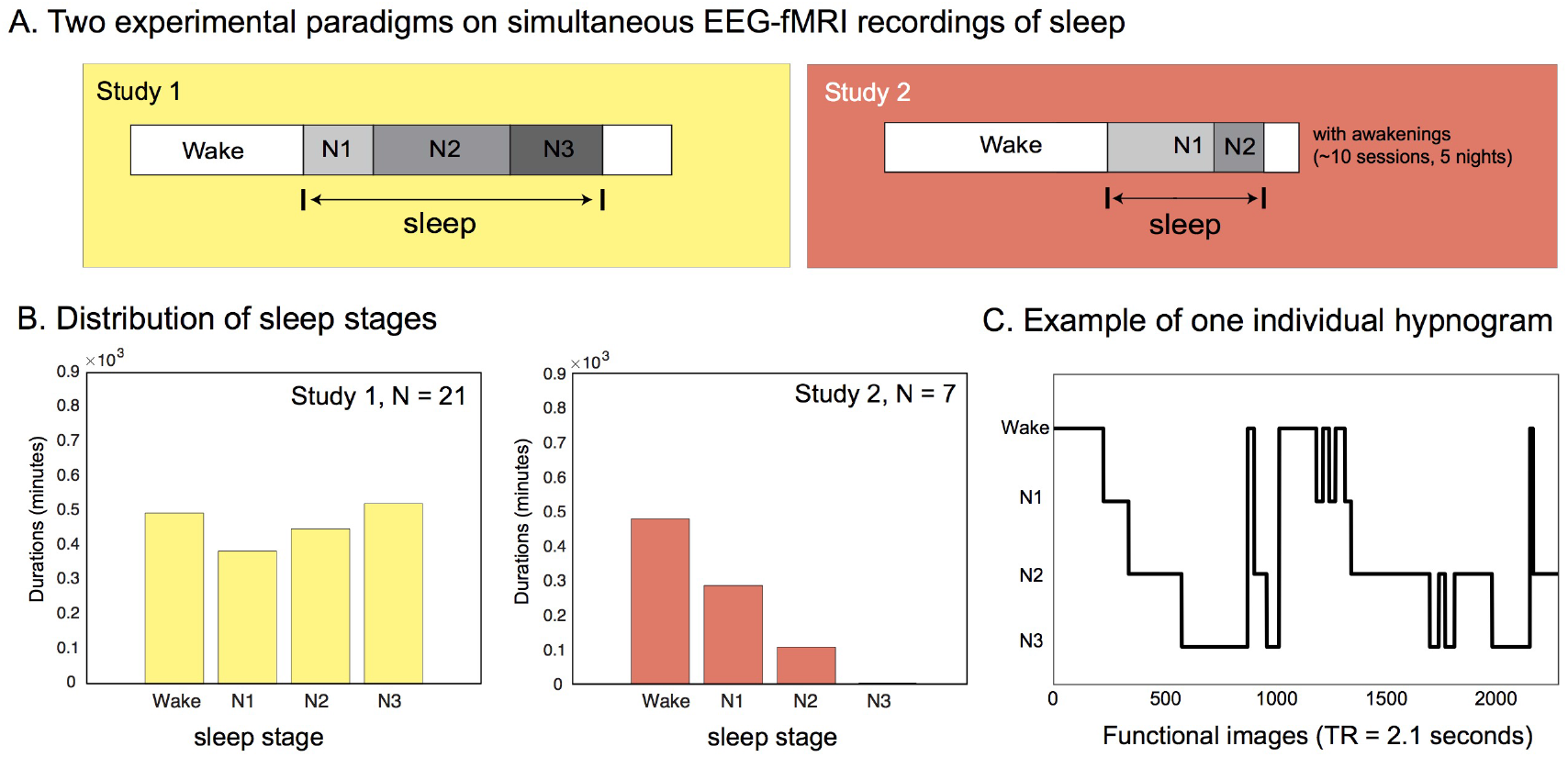
(A) The experiment for Study 1 lasted between 51 min to 2 hrs 40 min while Study 2 lasted between 5 minutes up to 20 minutes. (B) The distribution of accumulated data from each NREM sleep stage (in minutes of acquisition) shows that Study 1 covered up to N3, while Study 2 covered mostly wake and N1. (C) Example of sleep scoring (or hypnogram) for one participant in Study 1.

### 2.2 Spatial patterns in sleep and waking state

We applied a deconvolution process called total activation (TA) to all functional volumes of Study 1 and Study 2. *Transient frames* (*i.e.,* moments of activity changes) are extracted through a derivative step that is incorporated in the TA framework. Significant transient frames are then selected using a two-step thresholding process (see Supplementary Methods or (Karahanoğlu and Van De Ville, 2015)). Frames that survived are concatenated together and underwent a clustering procedure known as the iCAPs framework to obtain the most prevalent brain spatial patterns. We observed 17 large-scale brain networks displayed in Fig. 2, representing the different functional maps that dominate brain activity from wakefulness to deep sleep. The iCAPs are ordered in descending order according to the number of times that they appeared in the significant transient frames. We then looked at the wake/sleep stages where these significant transient points occurred, and identified which iCAP they corresponded to. Fig. 2 also shows the clustering distribution for each iCAP in pie charts, revealing the proportion of transient frames that the clustered iCAP occurred in each wake/sleep stage. Spatial similarities between iCAPs generated separately using Study 1 and Study 2 are shown in Supplementary Fig. SI 4.

**Fig. 2.**
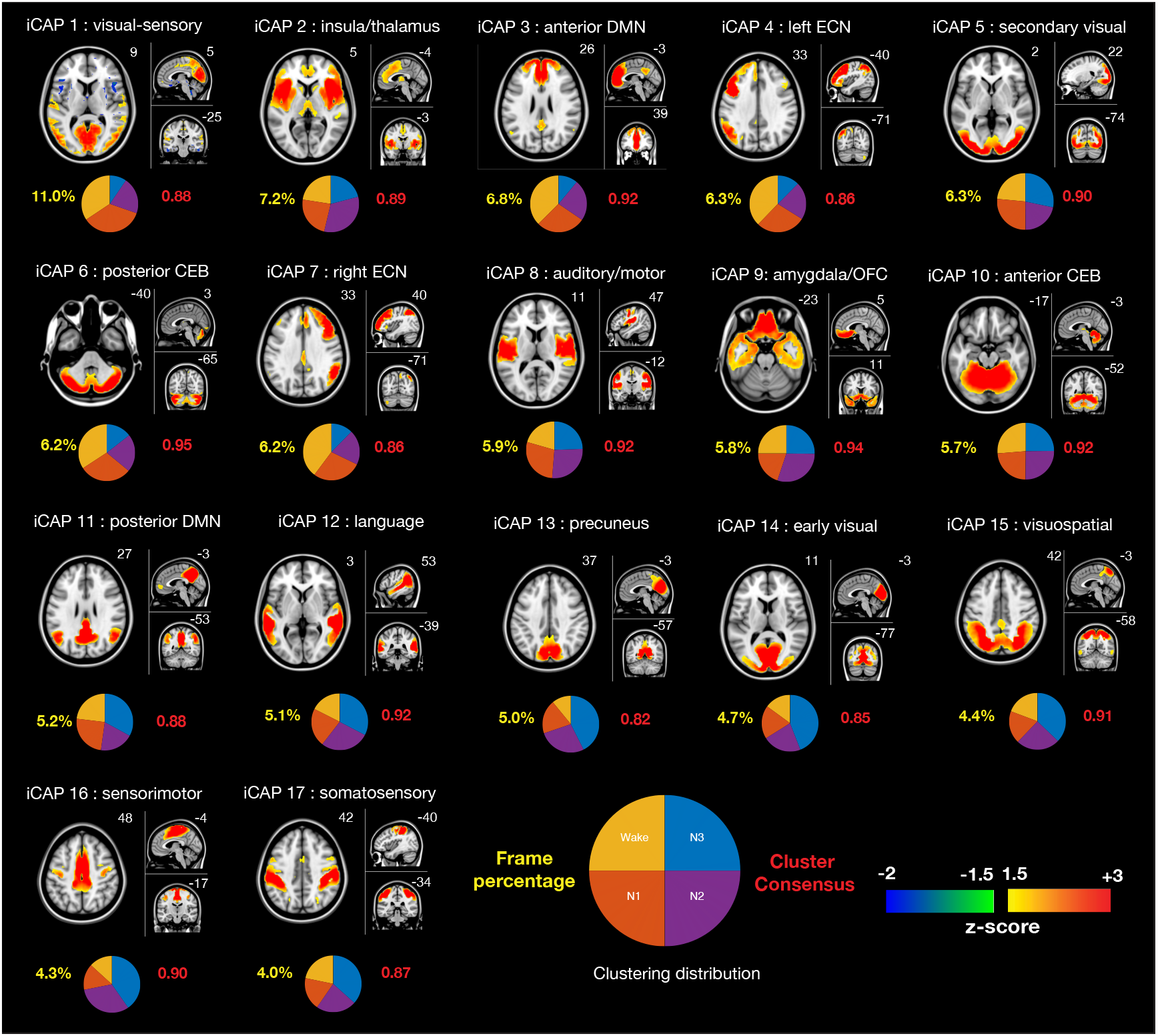
**Spatial patterns of the 17 innovation-driven co-activation patterns (iCAPs)** derived from all recordings across both studies. The iCAPs are numbered according to the percentage of significant transients that contributed to the recovery of that network (descending order), which are shown below the functional maps in yellow font. The cluster consensus of each iCAP is written in red. Pie charts indicate the distribution of each iCAP across sleep stage. MNI coordinates of each brain slice are indicated in white font. The names of the iCAPs are derived according to their correspondence with Greicius networks (Shirer et al., 2012) which are presented in the Supplementary Information (Table SI-1). CEB – cerebellum, DMN – default mode network, ECN – executive control network, OFC – orbitofrontal cortex.

The first iCAP (iCAP 1) featured both visual and somatosensory regions, and resembles a previously observed functional pattern that distinguishes sleep from waking conditions. These regions were found to be more prevalent in N1 and N2 sleep here, like in previous work (Tagliazucchi et al., 2013; Tagliazucchi and Laufs, 2014). Unlike any other iCAP extracted, this iCAP also reveals a negative activation in subcortical regions, very much similar to what was observed by Liu and colleagues (Liu et al., 2018), as is shown side-by-side in Fig. 3.

**Fig. 3:**
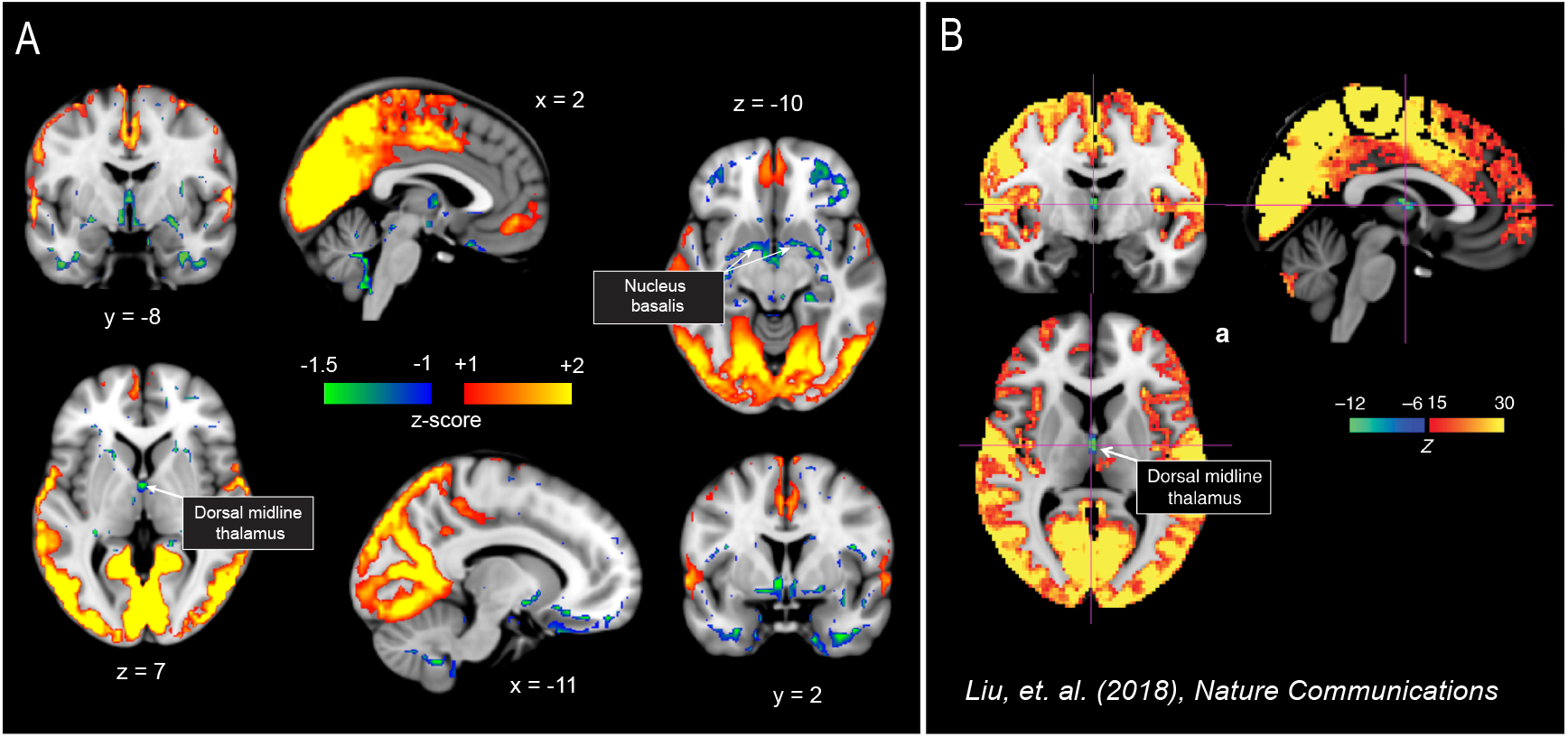
Visual-sensory iCAP reveals negative activations in subcortical regions. (A) Visual-sensory iCAP also shows negative patterns in subcortical regions. (B) Arousal-related network showing the deactivation in dorsal midline thalamus (figure adapted from previous study, Liu et al. 2018).

ICAP 2 featured regions of the salience network and presented a strong and selective activation of the insula, as well as portions of the thalamus. This network showed the highest likelihood to occur during N2 sleep. ICAP 3 predominantly occurred during wakefulness, displaying the anterior portion of the default mode network (DMN), and in particular the anterior cingulate. By contrast, iCAP 11 (posterior DMN) and iCAP 13 (precuneus, ventral DMN) were predominantly active during deep sleep (N3), and displayed stronger activation in the posterior regions (*e.g.,* PCC, precuneus, IPC). Notice that the anterior and posterior DMN were separately extracted during the clustering procedure despite the close similarity of their spatial patterns as shown in Fig. 4A. This reflects a strong dissociation between the DMN sub-networks in terms of their temporal dynamics across the different sleep stages.

**Fig. 4:**
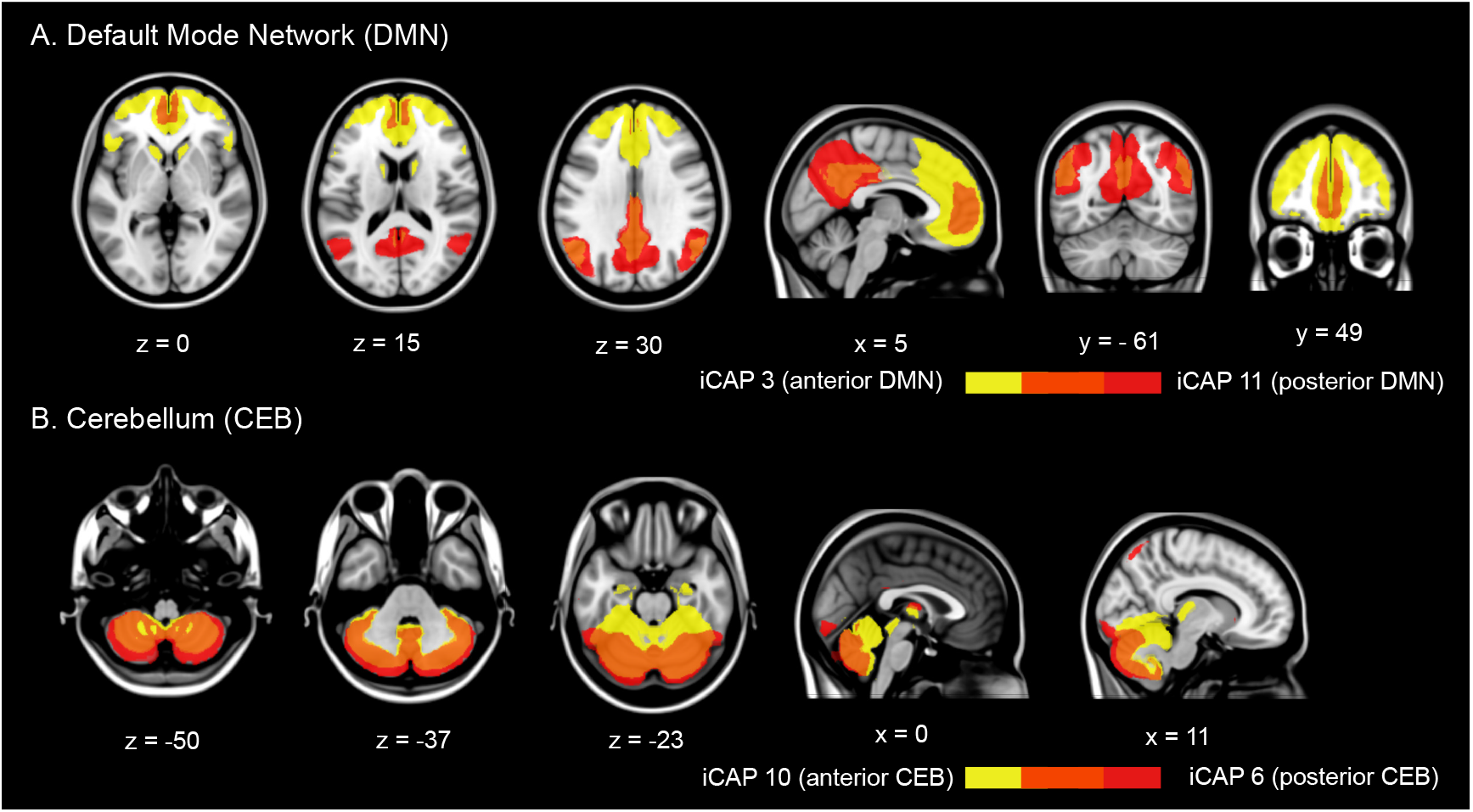
Dissociation of DMN and cerebellum into posterior and anterior parts. (A) Overlay of iCAPs corresponding to anterior DMN (yellow) and posterior DMN (red). (B) Overlay of anterior cerebellum (yellow) and posterior cerebellum (red). We also see some co-activation in the thalamus in the anterior cerebellum.

Meanwhile, attention-related networks such as the iCAPs 4 and 7 (left and right executive control (ECN) networks), and iCAP 15 (visuospatial) were also observed. Interestingly, both ECN networks mostly resulted from transients occurring during wakefulness, whereas the visuospatial network predominated during N3 sleep. ICAP 9 contained the amygdala and the orbitofrontal cortex, with limbic-emotional iCAPs predominating during N2. We also found various networks corresponding to sensory areas corresponding to transients from the deep sleep (N3). These included iCAP 8 (auditory/motor), iCAP 14 (early visual), iCAP 16 (sensorimotor), and iCAP 17 (somatosensory).

Finally, similar to the DMN, we also observed a dissociation of the cerebellum in the form of iCAPs 6 and 10, both featuring the anterior and posterior regions of the cerebellum, respectively. These iCAPs are overlaid in Fig. 4B. A detailed description of the regions in each iCAP (AAL regions (Tzourio-Mazoyer et al., 2002) and their similarity to Greicius networks (Shirer et al., 2012) are displayed in the Supplementary Information (Table SI-1).

### 2.3 iCAP relative cumulated durations are consistent with the cluster distribution of significant transients, and both reveal stage-dependent network activity

The iCAPs were obtained by clustering the significant transients from the data (i.e., instances when functional maps markedly increased or decreased activity). The pie charts in Fig. 2 display how the significant transients contributing to each iCAP were distributed across wake/sleep stages. In order to obtain the activity time-courses of iCAPs per participant in fMRI TR resolution (i.e., frame-wise), and be able to compute their durations (in contrast to only transients or activity changes), we performed a spatio-temporal transient-informed back-projection of the iCAPs onto the HRF-deconvolved frames.

In Fig. 5A, we display the relative cumulated durations (RCD) or the likelihood of each iCAP to appear in a specific sleep stage. This is a measure describing the cumulated time that a particular iCAP was active divided by the total time that the participant spent in a particular sleep stage. This normalization thus ensured that the reported persistence of an iCAP within one sleep stage was independent of the duration of that stage. Unsurprisingly, the relative cumulated iCAP durations appeared mostly consistent with the clustering distribution observed in Fig. 2. Attention-related iCAPs such as the left and right ECN showed predominant sustained activity in wakefulness. Conversely, sensory-related iCAPs (e.g., early visual, somatosensory, sensorimotor) were more persistent in N2 and N3 sleep compared to wakefulness and N1. The insula/thalamus iCAP also displayed predominant activity in N2 sleep. In addition, we found that the relative cumulated durations of iCAPs related to anterior DMN is higher during wakefulness, while the posterior DMN iCAP showed higher likelihood to appear in N3. These findings are well in line with previous observations of DMN dissociating into posterior and anterior parts upon reaching deep sleep (Larson-Prior et al., 2009; Sämann et al., 2011). Interestingly, we also observed the posterior (but not the anterior) cerebellum to be preferentially activated during wakefulness compared to N3. The corresponding test statistics (e.g., p-values, t-statistic, and effect sizes) for individual networks are reported in Supplementary Information Table SI-2.

**Fig. 5:**
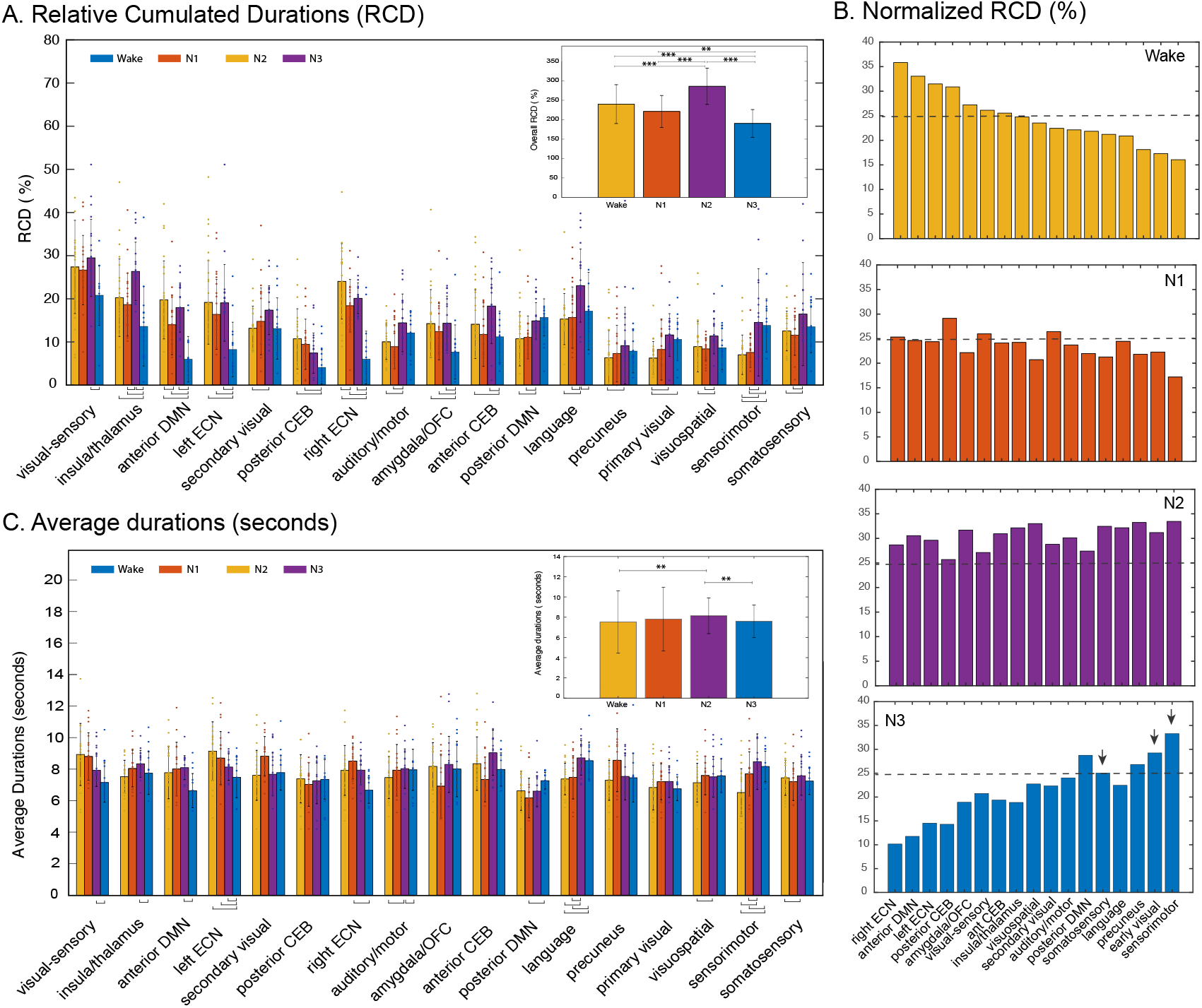
Duration of iCAPs in different sleep stages. (A) Relative cumulated durations (RCD, in %) of iCAPs depicting each network’s likelihood to occur in different sleep stages. The inset shows the overall trend of the iCAP durations in wakefulness/sleep stages, which are all above 100%, reflecting the tendency of iCAPs to overlap in time. (B) RCD divided by the number of time points that an iCAP is active in the whole time-course. The bars are ordered according to their descending representation in wakefulness. The broken horizontal line in each subplot indicates 25% likelihood, while the arrows in N3 subplot indicate visual and sensory-related iCAPs that are above the 25% line. (C) Average durations (in seconds) of iCAPs, reflecting the length of continuous activity of iCAPs. The inset corresponds to the overall trend of iCAP average durations. The height of the bars shows the mean and error bars indicate their corresponding standard deviations. The horizontal lines at the bottom of the bar plot correspond to significant differences evaluated through paired t-tests and permutation testing. Horizontal lines with 3 stars and 2 stars in the insets represent significant differences with p-values less than 0.001 and 0.01, respectively.

We then looked at the general trend of network activity across the different sleep stages (inset of Fig. 5A), and found that the RCD of iCAPs significantly increased during N2 with respect to wakefulness and N1, followed by a steep decrease in N3. Please note that values above 100% are due to the temporal overlapping nature of iCAPs, with two or more iCAPs occurring at the same time. We then computed a network-based normalization of RCD, displayed in Fig. 5B. This is equivalent to normalizing the number of active time-points in each sleep stage by successive division to two factors: (1) overall number of time instances that an iCAP is present over the whole duration of the recording and (2) the total time that the participant spent in a particular sleep stage.

The bar plots in Fig. 5B show that almost all iCAPs displayed proportions above 25% in N2 sleep (broken horizontal line). We observed a marked disparity in iCAP proportions during wakefulness and N3, in contrast to the more uniform distribution in N1 and N2 sleep. Moreover, iCAPs that were more represented in wakefulness were less represented in N3, which is consistent with the RCD measures of each iCAP across wake/sleep stages in Fig 5A (all p<0.05). Specifically, iCAPs that reached more than 25% activity during specific sleep/wake stages are: (1) wakefulness -- visual-sensory, left and right ECN and posterior CEB, (2) N1 --posterior CEB, visual-sensory, secondary visual, and (3) N3 -- posterior DMN, somatosensory, precuneus, early visual, sensorimotor.

Contrasting with the RCD above, the average duration or mean lifetimes of an iCAP represents the average bouts (in seconds) of continuous activity. Overall, we observed iCAPs to be active between 5 to 10 seconds (7.3 ± 1.7 seconds; Fig. 5C). In general, we also found iCAP activity to have longer durations in N2 compared to N3 (p < 0.01), and compared to wakefulness (p < 0.01). For some of the networks, the average durations were proportional to their relative propensity to occur in each sleep stage. For instance, the left ECN displayed the least likelihood to occur in N3, and also exhibited the shortest average duration during that stage.

### 2.4 Alterations in network couplings in different sleep stages

To go beyond the iCAPs’ individual temporal properties, we looked at the probability of having either none, one, two, or more iCAPs occurring at one time-point. Fig. 6A shows the likelihood for different numbers of overlapping iCAPs to occur in wakefulness and in different sleep stages. We observed the overlaps to be maximally N = 10, and thus we limit the histogram for the range {N = 0,1,2, …, 10}. Among all vigilance states, N2 displayed a relatively flatter distribution especially compared to wakefulness and N3 (both p < 0.05), with a higher likelihood of having 2 to 5 overlapping iCAPs. For a table of test-statistics, see Supplementary Table 3. Given that individual iCAPs had higher likelihood to occur in N2 compared to other sleep stages, it is not surprising that iCAPs are also more likely to overlap in time during N2.

**Fig. 6:**
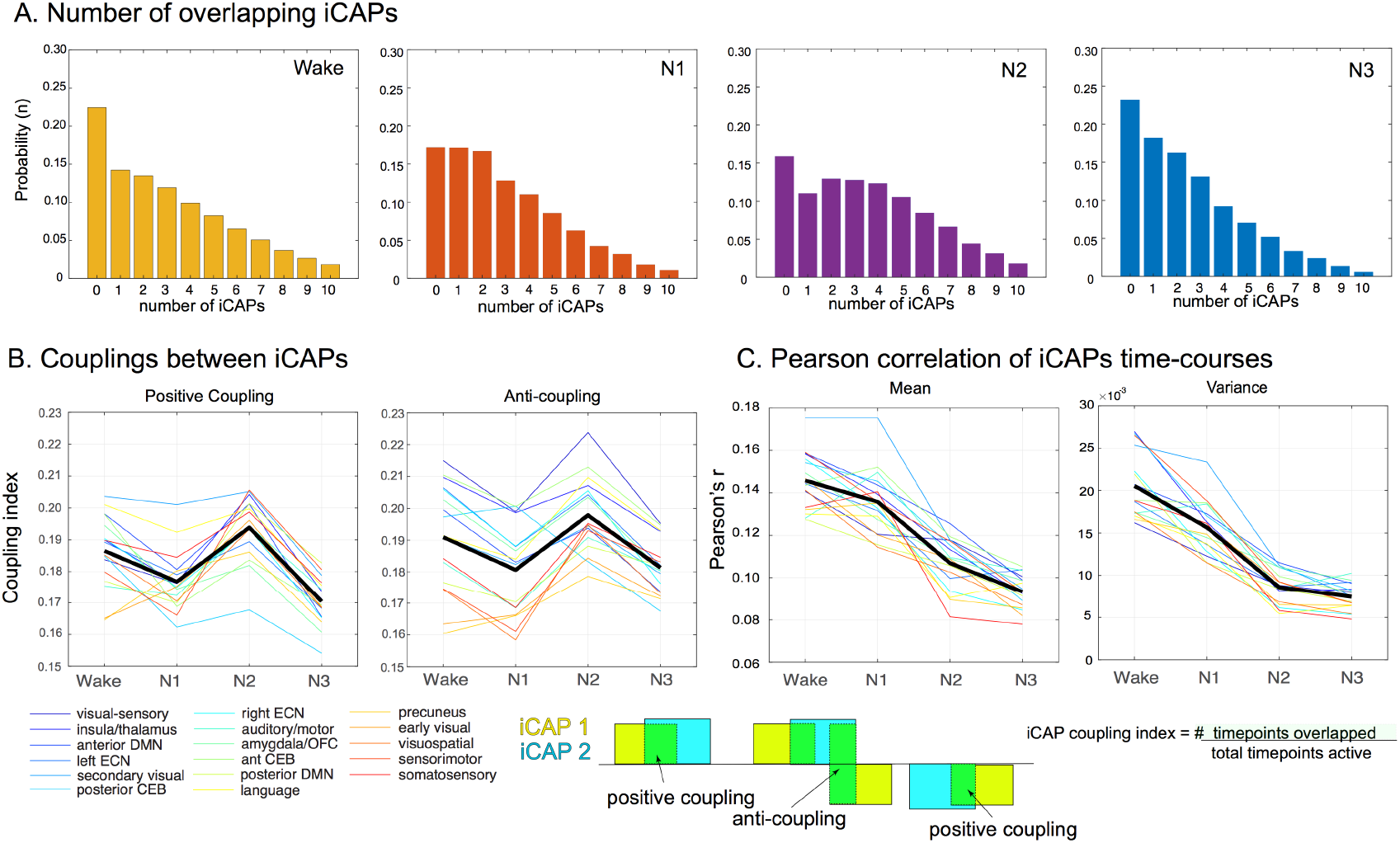
Interactions between iCAPs across different sleep stages. (A) The probability of different numbers of iCAPs to overlap is displayed for wakefulness and different sleep stages. (B) The iCAP coupling index pertains to the number of time-points during which a pair of iCAPs were both active, divided by the total number of time-points that at least one of them was active. This was computed separately for similar signed activations (same-signed coupling) or opposite signed activations (opposite-signed coupling). Coupling was computed for each network (i.e., pair-wise coupling of one network with all the other networks). (C) Mean and variance of classical FC metric (Pearson’s r) applied to iCAP time-courses. The computation for the coupling index between pairs of iCAP is illustrated on the bottom of the figure.

We then evaluated the pair-wise coupling between each iCAP and all other networks (Fig. 6B). The coupling index is a normalized measure of the number of time-points at which an overlap occurs between two pairs of iCAPs, divided by the total number of time-points that at least either one is active. This is different from the temporal overlap measure observed in Fig. 6A because we focus on pair-wise coupling between two iCAPs. This can also be referred to as the Jaccard similarity score of two iCAP time-courses. We also take into account the signs of the activations, see illustrations in Fig 6. In general, all iCAPs showed a high mean same-signed coupling and opposite-signed coupling in wakefulness and N2, and a decrease in N1 and N3 (p<0.001, see Supplementary Table 4). Next, we compared these observations using classical FC analysis (i.e., Pearson correlation) applied to the iCAP time-courses. We observed that the pairwise network FC across iCAPs decreased with increasing sleep depth (with no increase in N2). This observation is consistent with the general notion that connectivity breaks down with increasing sleep depth (Tagliazucchi and Laufs, 2014). The variance observed in Fig. 6C reveals low disparity of correlation between iCAP time-courses during N2 and N3 sleep, about 3 times less than what is observed in wakefulness and N1.

## 3. Discussion

### 3.1 General findings

Here, we extracted large-scale brain networks or iCAPs by clustering moments of significantly changing brain activity from two studies using simultaneous EEG and fMRI recordings during sleep. The use of transient fMRI activity allowed us to obtain temporally overlapping spatial patterns together with subject-specific time-courses at a time-scale of seconds. We found that coordinated activation of regional brain areas during wakefulness, or iCAPs, was largely preserved in N1, N2 and N3 sleep, yet with specific relative distributions and dynamic modulations across different stages of wakefulness and sleep. This first observation is largely consistent with previous work reporting the persistence of large-scale brain networks during sleep (Horovitz et al., 2009; Sämann et al., 2011), with various modifications when reaching deeper sleep stages, such as decreased connection strengths in RSNs (Larson-Prior et al., 2009), changes of hierarchical organization into smaller independent modules (Boly et al., 2008), or reduced long-range temporal dependencies in the BOLD signal (Tagliazucchi et al., 2013). For instance, the well-known attention-related networks, such as the bilateral ECN, occurred preferentially during wakefulness and N1, whereas networks associated to primary sensory systems were most present during N2 and N3, consistent with some earlier observations (Horovitz et al., 2008; Larson-Prior et al., 2009). Whenever active, these networks sustained continuous bouts of activity on an average of about 7.3 ± 1.7 seconds across all subjects and all sleep stages. For some networks, their mean lifetimes changed as a function of sleep stages. Furthermore, pair-wise iCAP couplings generally show marked increase in N2 sleep followed by a decrease in N3. This pattern is also manifested in terms of the observed tendency of networks to overlap the most during N2 sleep.

In the next subsections, we discuss the most important findings of the work. We also give interpretations to the observed dynamical characteristics of cross-network interactions and their relation to the integrated information theory (IIT) of consciousness (Tononi, 2004).

### 3.2 Visual-sensory iCAP confirms the emergence of arousal-related network

The first iCAP (iCAP 1, Fig. 2), corresponding to the network of regions contributing the highest number of significant transients in our dataset, was the visual-sensory iCAP, which predominated during the transition to sleep (N1). It included a large portion of the visual cortex, as well as some sensory and motor regions. This network also displayed a unique characteristic of having negative activations in subcortical areas. Using combined electrophysiological and fMRI signals, Liu and colleagues (2018) described a very similar spatial pattern, with negative activation in basal forebrain and thalamus paralleled by positive activations in sensory cortices, which arose during momentary drops in arousal, indexed by a spectral shift in local field potentials toward low frequencies. In Fig. 3, we provide a side-by-side comparison between our visual-sensory iCAP 1 and the arousal-related network observed by Liu et al. While both networks displayed positive activations in visual, sensory and motor regions, they also exhibited negative activation in the dorsal midline thalamus, as well as the nucleus basalis. Because the thalamus holds an important role in regulating the physiological state of arousal during sleep (Gent et al., 2018; Saper et al., 2010) and is responsible for relaying motor and sensory signals to the cerebral cortex, we can suggest that reduced thalamic activity favors sleep onset while limiting incoming external sensory stimulation. This interpretation is also supported by the fact that most of the frames that contributed to this iCAP came from wake and N1, suggesting that this iCAP might play a key role in the onset of sleep. Interestingly, however, we found that this network exhibited a persistent likelihood in terms of the cumulated durations to occur in N2, before decreasing in N3. Fittingly, it has also been found in previous studies that there is an increase in BOLD signal variance in sensory and motor cortices during N1 and N2 sleep, which has been used as diagnostic for detecting sleep in typical resting-state recordings. On the other hand, high activities in frontal, parietal, and temporal cortices are typically associated to wake conditions (Tagliazucchi et al., 2013; Tagliazucchi and Laufs, 2014). These two brain spatial patterns bear resemblance to the visual-sensory iCAP and the bilateral ECN, respectively, whose temporal properties show consistent behavioral stage-dependence. In addition, Stevner et al. (2019) reported the same visual-sensory network to be implicated also in N1.

### 3.3 DMN and cerebellum dissociates into posterior and anterior regions

The DMN has been well studied particularly in light of its connectivity changes when subjects transition from wake to light sleep, and deep sleep. The initial hypothesis was that the DMN supports a range of self-related mental processes, such as unconstrained self-referential thought and recollection. However, previous studies have found the DMN to persist during light sleep, as well as in deep sleep with reduced connectivity between the medial prefrontal cortex (MPFC) and the rest of the network (Horovitz et al., 2009; Larson-Prior et al., 2009). In the present work, we observed a dissociation of the DMN sub-components into its posterior and anterior parts (Fig. 4A). The pie-chart distributions of cluster frame indices displayed in Fig. 2 and the cumulated durations in Fig. 5A reveal that the anterior region of the DMN mostly occurred during wake and N1 sleep whereas the posterior DMN predominated in N2 and N3 sleep. We found a similar antero-posterior dissociation in the cerebellum (overlap shown in Fig. 4B). Studies that investigated FC in the posterior and the anterior regions of the cerebellum have remained inconclusive regarding their potential roles in sleep. The cerebellum has been found to be related to motor control and motor memory formation (Gao et al., 2012). It has also been observed to show sleep stage-dependent activity, whose impairments disrupt the sleep-wake cycle which then leads to sleep disorders (Del Rosso, L.M, Hoque, 2014). As a general observation, signals in the cerebellum appear to be lower during N1 as compared to wakefulness (Hiroki et al., 2005; Kaufmann et al., 2006). However, a map summarizing major cerebellar research in sleep using fMRI and PET revealed a localized change in cerebellar activity during SWS surrounding the cerebellum’s larger lobules (parts IV, V, VI, and VII), as well as a marked correlations with slow spindles in N2 (Canto et al., 2017). These lobules make up the anterior region of the cerebellum (Dang-Vu et al., 2008). Moreover, it has been previously shown that the sensorimotor domain is related to the anterior cerebellar lobe, while the cognitive domain corresponds to the posterior cerebellar region (Stoodley and Schmahmann, 2010). This particular finding is consistent with the observed co-activation of the anterior cerebellum with regions corresponding to the sensorimotor network represented by iCAP 10 (see also Shirer et al. 2012) and Supplementary Table SI-1) Meanwhile, the inferior posterior lobe has been associated to the frontal gray matter, a well-known core area of executive function (Jung et al., 2019; Tiemeier et al., 2010). Altogether, these studies support our current findings on the preferential tendency of posterior and anterior cerebellum to persist in wakefulness and N2, respectively.

### 3.4 Network temporal characteristics in sleep and their relation to the information integration theory of consciousness

Previous connectivity analyses of brain regions found a general decrease of FC with increasing sleep depth (Haimovici et al., 2017; Spoormaker et al., 2010; Tagliazucchi et al., 2016). Classical FC approaches capture statistical inter-dependencies between activity from two brain regions (Friston, 2011). Typically, if two regional time-courses exhibit simultaneous positive (or negative) values over some time-windows, and exactly opposite values on some other time-windows, the final correlation (and derived FC) between both regions would be close to zero, despite the synchronized deviations from the baseline, albeit the incoherent signs. Therefore, it follows that a decrease in FC does not mean a decrease in global network activity. This clarification is particularly important with regards to the interpretation of Tononi’s information integration theory of consciousness (2004) which has been cited by several neuroimaging studies that show marked reduction of brain network integrity (Nofzinger, 2006) from wakefulness to deep sleep.

In this work, we instead demonstrated that network activations increased when participants went from wakefulness to N2, while they decreased when reaching N3. This important observation was captured because we were able to extract network temporal activity at the frame-wise level for each individual iCAP. In contrast, classical dynamic FC analyses only capture the statistical relationship among ROIs comprising the networks, and not the actual activity of the networks themselves. Furthermore, because the iCAP approach allows networks to overlap in time, we were able to observe that functional networks show higher likelihood to overlap (e.g., 2 to 4 iCAPs active at the same time) in N2 compared to other stages. Congruently, we also found high coupling indices in N2. What is remarkable is the simultaneous increase in both the same-signed and opposite-signed activations in N2 (Fig. 6B). By contrast, N1 and N3 generally showed much lower coupling indices. Meanwhile, classical Pearson correlation measure applied to iCAP time-courses displayed a decreasing trend from wakefulness to deep sleep, consistent with FC findings based on regional fMRI time-courses (Haimovici et al., 2017; Kung et al., 2019; Spoormaker et al., 2010; Tagliazucchi et al., 2016).

We therefore summarized three important findings regarding network temporal characteristics of sleep stage N2: (1) FC between networks decreases with sleep depth, (2) there is a higher likelihood for stereotyped networks or iCAPs to emerge in N2 compared to wakefulness and other sleep stages, and (3) both same-signed and opposite-signed coupling indices between iCAPs increased in N2. We would like to speculate that decreased FC reflects an overall reduced efficiency of information transfer between networks, well in-line with the established concomitant reduction of consciousness (Tononi, 2004) during sleep. Meanwhile, the increased network activity in N2 manifests unstable cross-network talk. In other words, observation (1) would correspond to a decrease in global network integration, while observations (2) and (3) express network attempts to communicate specifically during N2, which would result in increased inter-regional interactions with nonetheless unstable synchronizations. This finding is in agreement with a very recent observation by Kung et al (2019), who reported a very high variance in inter-network mean dynamic FC across sliding windows during N2, compared to during wakefulness, N1 and N3, while mean dynamic FC decreased from wakefulness to N3. High dFC variance in N2 was interpreted as elevated instability of information transfer between networks, whereas low mean dFC would reflect decreased intra-network consistency.

Concerning N3, we observed (1) an overall decrease of global network occurrence, (2) a decrease in network mean lifetimes, and (3) a decrease in functional association characterized by lowest cross-network FC. Deep sleep is first and foremost characterized by unresponsiveness, and the decline of ability to react to external stimuli (Cirelli and Tononi, 2008). Altogether, our results are consistent with a more stable brain state (Jobst et al., 2017) yet more localized signal integration in SWS (Deco et al., 2017). Spatially, the huge drop in anterior DMN’s likelihood to occur in deep sleep compared to other stages, together with the high persistence of primary sensory networks, provide further support to the notion of a break-down of long-distance functional connections (Tagliazucchi et al., 2013) in favor of short-distance associations (Boly et al., 2012). The global decrease of network activity and mean lifetimes in N3 is an indicator of a more stable state of the system, but also a general loss of effective information integration.

### 3.5 Methodological Aspects

The iCAPs framework has already been applied to resting-state fMRI data of healthy (Karahanoğlu and Van De Ville, 2015) and clinical populations (Zöller et al., 2019). The recovered iCAPs in the present study were highly similar to the ones observed in these two previous studies, while additional spatial patterns unique in the sleep were also found (e.g., visual-sensory network with deactivations in subcortical regions, cerebellum, and insula). Moreover, the recovery of both anterior and posterior DMN confirms previous findings on the dissociation of DMN with increasing sleep depth, thereby further evidencing the framework’s performance at extracting coherent large-scale brain activity. The framework is unique in its ability to detect transients which allows for the extraction of spatially and temporally overlapping functional networks, a particular advantage that is beyond the classical methodologies applied in sleep studies. Using this advantage, we were able to show, for the first time, individual network activity at precise fMRI temporal resolution across sleep stages and found distinct stage-dependent activity for each network.

### 3.6 Summary

In summary, we investigated the dynamic properties of large-scale functional networks during wakefulness and NREM sleep. We used the TA and iCAPs framework to capture precise moments of transient brain activity, giving us a quantitative view of how the brain dynamically evolves across the different stages of NREM sleep. We found new networks that are largely related to regions that support the physiological organization of sleep and arousal. We also uncovered whole brain spatial patterns that resemble currently known RSNs whose temporal profiles are consistent with previous findings. The temporal dynamics these networks exhibited alterations between different sleep stages. In particular, we observed a global dissociation/decrease of brain activity both in spatial and temporal domains, from wakefulness to deep sleep. Unexpectedly, we found an increase in network activity in N2 and a global increase in simultaneous positive and negative network co-occurrence, signaling instability of network synchronization and ineffective brain integration. Altogether, these findings support the information integration theory of consciousness during sleep and provide new evidence for the presence of unstable yet distributed global inter-regional interaction in N2 sleep.

## 4. Materials and Methods

### 4.1 Functional MRI datasets

We used two complementary sets of simultaneously recorded EEG-fMRI data acquired in the context of two different studies. Ethics approval was obtained for each study and all participants signed written informed consent forms. The first dataset came from a study on sustained sleep (denoted from here onwards as *Study 1*), where participants were allowed to sleep for as long as they could manage, while EEG and fMRI data were simultaneously and continuously recorded. Twenty-six right-handed healthy participants participated in this study. All of the participants filled out questionnaires and underwent a semi-structured interview that established the absence of neurological, psychiatric, or sleep disorders. They were non-smokers, moderate caffeine drinkers, and were not taking any medication. The experiment was conducted at around 10 pm and the total recording time lasted between 51 min and 2 hrs 40 min (mean: 1 hr 43 min). We excluded a total of 5 participants who were not able to sleep inside the scanner (resulting in 21 subjects, mean age ± SD: 22 ±2.4 years; 15 women).

The second dataset focused on the sleep onset period (denoted below as *Study 2*). Ten healthy subjects (4 female) underwent simultaneous EEG-fMRI recording. All participants had no history of neurological or psychiatric disease, no previous or current use of psychoactive drugs, and were non-smokers. The experiment was conducted at around 9:30 pm and the awakening recording lasted 1 hr 30 mins. Participants were placed in the MRI scanner and asked to fall asleep. Successive awakenings of up to 10 rounds per session were performed during this period to record possible dreaming that occurred. The whole experiment included around 4 to 5 sessions (5 nights) per participant. As expected from this experimental design, in the majority of the recordings, participants were in the awake state to light sleep (N1). Fig. 1 displays the two experimental paradigms, together with the distribution of sleep states reached by participants. We excluded 3 subjects due to difficulty in scoring the EEG data (7 subjects remaining, mean age ± SD: 20.5 ± 1 year; 5 women). In the present work, we did not investigate the dream reports collected in Study 2 because our main goal was to characterize dynamic neurophysiological changes occurring from wakefulness to deep sleep, and also because we did not collect dream data in Study 1 (waking up participants would have prevented them from reaching deeper sleep stages).

### 4.2 EEG preprocessing and sleep scoring

For the data in Study 1, the EEG setup included a 64-channel MRI-compatible EEG cap, two pairs of electrocardiogram (ECG), horizontal and vertical electrooculography (EOG), and 1 pair of chin electromyographic (EMG) electrodes (BrainAmp MR plus, Brain Products GmbH, Gilching, Germany). For the data in Study 2, a 14-channel MR-compatible system (Brain Products GmbH, Gilching, Germany) was used, plus a total of ten cortical (EEG) electrodes. Two of the ten cortical EEG electrodes served to measure EOG, three EMG chin electrodes, and one ECG electrode on the back were used to monitor participants’ sleep patterns. The preprocessing of both datasets was done using the Brain Analyzer software (Brain Products GmbH, Gilching, Germany). Specifically, gradient artifacts were removed offline using a sliding average of 21 averages and subsequently, the EEG data, initially sampled at 5000 Hz, was down-sampled to 500 Hz and low-pass filtered with a finite-impulse response filter with a bandwidth of 70 Hz. For the 64-channel recordings in Study 1, ICA was the most reliable method to remove ballistocardiogram and oculo-motor artifact, while a template subtraction method (Allen et al., 1998) was used for the 14-electrode recordings in Study 2.

The preprocessed EEG data for Study 1 and Study 2 were then scored by three experts each, according to standardized American Academy of Sleep Medicine (AASM) polysomnographic criteria for sleep scoring (Iber, 2007).

### 4.3 Functional MRI acquisition and preprocessing

For Study 1, MRI data were acquired on a 3 Tesla whole-body MR scanner (Tim Trio, Siemens, Erlangen, Germany) using a 12-channel head coil. Functional images were acquired with a gradient-echo EPI sequence (repetition time [TR]/ echo time [TE]/flip angle = 2100 ms/40 ms/90) and parallel imaging (GRAPPA; acceleration factor = 2). Each functional image comprised 32 axial slices (thickness = 3.2 mm without gap, FOV = 235 × 235 mm, matrix size = 128 × 84, voxel size: 3.2 × 3.2 × 3.84 mm,) oriented parallel to the inferior edge of the occipital and temporal lobes. On average 2789 functional images were recorded during one continuous scanning session (between 1459 and 3589 images). Structural images were acquired with a T1-weighted 3D sequence (MPRAGE, TR/inversion time [TI]/TE/flip angle = 1900 ms/900 ms/2.32 ms/ 9, FOV = 230 × 230 × 173 mm3, matrix size = 256 × 246 × 192 voxels, voxel size = 0.9 mm isotropic).

For Study 2, MRI recording was performed on a 3 Tesla Philips Achieva MRI scanner. Functional images were acquired using a gradient-echo EPI sequence (repetition time [TR]/ echo time [TE]/ flip angle = 3000 ms/ 30 ms/ 83). Each functional image comprised 50 axial slices (thickness = 2.5 mm without gap, FOV = 96 mm × 96 mm, voxel size: 2.5 mm isotropic) oriented parallel to the inferior edge of the occipital and temporal lobes. Structural images were acquired with a T1-weighted 3D sequence (TR/inversion time [TI]/TE/flip angle = 1570 ms/8.4 ms/3.42 ms/ 8, FOV = 256 × 256 × 220 mm3, matrix size = 256 × 256 × 220 voxels, voxel size = 0.929 mm × 0.929 mm × 1mm).

Both functional datasets were first preprocessed following standard procedures (Van Dijk et al., 2010). Functional volumes were realigned to their mean images using SPM12 (http://www.fil.ion.ucl.ac.uk/spm/). The realigned functional images underwent nuisance regression (mean white matter and cerebrospinal fluid, constant, linear, and quadratic drifts). We then applied spatial smoothing using an isotropic Gaussian kernel of 6 mm full width at half maximum. For the final analyses, the first 10 volumes were discarded to achieve steady-state magnetization of the fMRI data. To correct for large motion, we applied scrubbing (Power et al., 2012) in the functional data by marking time-points with framewise displacements of more than 0.5 mm. The iCAPs framework require a constant sampling rate, therefore, marked frames were replaced by interpolating the disconnected frames using spline interpolation (Karahanoğlu et al., 2013). Finally, structural scans (T1) were co-registered to the mean functional image, and the transformed volume was segmented using SPM 12 Segmentation to obtain probabilistic gray matter masks.

### 4.4 Total Activation and iCAPs framework

We applied the Total Activation (TA) procedure to the subject-space fMRI recordings for each participant of Study 1 and Study 2 datasets. The TA procedure started by a deconvolution of fMRI time-series at each voxel to remove hemodynamic effects (Farouj et al., 2017; Karahanoğlu et al., 2013). Temporal transitions or *transients* (changes in activity) were then derived for each participant in Study 1 and each session of each night in Study 2. Highly significant transient frames (functional images) were identified and normalized to MNI coordinate space. The normalized functional images of all subjects were then concatenated, forming one single data matrix with all significant transients. The data matrix with a dimension of number of voxels times number of significant transients was fed into a clustering procedure to obtain temporally co-activating brain patterns. Because the functional networks were obtained by clustering significant transient brain activity, also known herewith as innovations, the resulting co-activation patterns are referred to as *innovation*-*driven* co-activation patterns, hence the name *iCAPs*. The optimal number of clusters used in the clustering procedure was determined using consensus clustering (Monti et al., 2003), which implements a multiple subsampling procedure. We evaluated multiple numbers of cluster values in the set K = {10, 11, 12, …, 25}. The optimal K was selected based on different stability metrics. One measure is the *cluster consensus* which reflects the consistency of a transient frame to be clustered in the same iCAP. After obtaining stable iCAPs, we used a spatio-temporal transient-informed regression (Zoller et al., 2019) to extract the activity time-courses of iCAPs for each participant. The TA and iCAPs pipelines are publicly available and additional information as well as the mathematical details of the whole method are laid out in the Supplementary Methods.

### 4.5 Extraction of temporal characteristics and network interactions in each sleep stage

Comparison of the iCAPs time-courses across different sleep stages was performed by z-scoring the entire iCAP time-courses per participant’s recording, and thresholding at |z| > 1.5. The time points that survived thresholding were considered as “active”. The choice of threshold was motivated by previous works that implemented TA and iCAPs framework (Karahanoğlu and Van De Ville, 2015; Zöller et al., 2019). We then computed the temporal characteristics (e.g., cumulated durations, average durations, temporal overlaps, and network interactions or *couplings*) of iCAPs in each sleep stage. Scrubbed frames (*i.e.,* those with framewise displacement more than 0.5 mm) are taken into account and are not included in each temporal measure explored. The relative cumulated durations (RCD) computed in percentage pertains to the likelihood of an iCAP to occur in wakefulness and in each sleep stage. This is computed by counting the number of time-points that an iCAP is active divided by the total length of time a participant spent in that particular sleep stage. The overall RCD can go above 100% due to the overlapping nature of the iCAP time-courses (*i.e.,* more than one iCAP can occur at one time-point). The average duration, measured in seconds, is the length of time that an iCAP is continuously active. We also evaluated the interaction of iCAPs with other networks by computing the overall percentage of temporal overlaps in wakefulness and sleep, as well as the number of pair-wise iCAP co-activations. The latter is represented by the coupling index which reflects the number of time-points during which a pair of iCAPs were both active divided by the total number of time-points that at least either one of them was active. We take into account the signs of the activations (similarly signed or opposite signed). Z-scoring the time-courses allows positive and negative activation signs. Positive (negative) activation signs reflect the time-points when an iCAP is at least 1.5 standard deviations above (below) the overall mean amplitude of iCAPs.

### 4.6 Statistical Analysis

Duration measures of individual iCAPs were compared across the different sleep stages using paired t-tests and 1000 rounds of permutation testing (randomly shuffled). The corresponding t-statistics, p-values and effect sizes are displayed in Supplementary Table 2. The overall comparison of iCAP cumulated durations (general trend), network temporal overlap and coupling indices were done through the Analysis of Variance (ANOVA), and the p-values were obtained using successive multiple comparison test (Tukey’s range test). The corresponding F-statistics (ANOVA) and the corrected p-values for the temporal overlap and coupling indices are displayed in Supplementary Table 3 and 4, respectively. For the analysis of temporal characteristics (e.g., average durations, RCD, temporal overlaps, and couplings), we only included subjects who reached N3, and thus we only limit this part of the analysis to Study 1.

## 5. Acknowledgements

This work was supported by the Swiss National Science Foundation (Grant number 205321-163376) to A.T.; the National Institute for the Humanities and Social Sciences (NIHSS, Grant number SDS14/1117) to D.A.W.; the German Academic Exchange Service (DAAD, grant number 91570813) to D.A.W.; the National Research Foundation of South Africa (NRF, grant number 83341) to D.A.W.; the Harry Oppenheimer Memorial Trust, South Africa (OMT grant number 20027) to D.A.W; the National Center of Competence in Research (NCCR) Affective Sciences financed by the Swiss National Science Foundation (grant number, 51NF40-104897) and hosted by the University of Geneva; the Swiss National Science Foundation (grant numbers: 320030-159862, 320030-135653) to S.S.; and the Mercier Foundation.

We would also like to acknowledge all participants in the two experimental studies included in this work, and Carina Zoellner and Annika Rips for help with data acquisition in Study 2.

## 6. Author Contributions

Conceptualization, A.T. and D.V.D.; Methodology, A.T. and D.V.D.; Validation, A.T.; Formal Analysis, A.T.; Investigation, D.W.A. and V.S.; Resources, D.W.A., V.S., L.P., L.B. and M.S.; Data curation, A.T., D.W.A., V.S., L.P., and L.B.; Writing – Original Draft, A.T.; Writing – Review & Editing, A.T., D.W.A., L.P., N.A., S.O.S. and D.V.D.; Visualization, A.T.; Supervision; S.O.S. and D.V.D.; Funding Acquisition, M.S., S.O.S., and D.V.D.

