## Supplementary Information for "NREM sleep stages specifically alter dynamical integration of large-scale brain networks"

### Supplementary Information for the article "*NREM sleep stages specifically alter dynamical integration of large-scale brain networks*"

#### Supplementary Methods

*Deconvolution of fMRI signal via Total Activation*

The total activation (TA) framework is the process of deconvolving fMRI time series to detect time points where there are significant changes in brain activity (Farouj et al., 2017; Karahanoğlu et al., 2013). Fig. SI 1 illustrates the TA framework. At each voxel $i$, fMRI signals $y(i,t)$ are modelled as a block-like activation activity-inducing signal $u(i,t)$ convolved with the hemodynamic response function $h(t)$, and corrupted by a Gaussian noise $\varepsilon(i,t)$:

$$y(i,t) = h(t) * u(i,t) + \varepsilon(i,t)$$

The denoised fMRI signal, also called herewith as activity-related signal $x(i,t)$ (or in matrix form $\boldsymbol{X}$), is obtained by running a combined spatial and temporal regularization that aims to minimize the following cost function:

$$\widetilde{\boldsymbol{X}} = argmin \frac{1}{2} ||\boldsymbol{Y} - \boldsymbol{X}||_F^2 + R_T(\boldsymbol{X}) + R_S(\boldsymbol{X})$$

The temporal and spatial regularization are given by the following terms:

$$R_T(\boldsymbol{X}) = \sum_{i=1}^{N_i} \lambda_T(i) \sum_{t=1}^{N_t} |\Delta\{\boldsymbol{X}(i,.)\}|$$

$$R_S(\boldsymbol{X}) = \sum_{t=1}^{N_t} \sum_{i=1}^{N_i} \sqrt{\sum\nolimits_{i \in S} \Delta_{Lap}\{\boldsymbol{X}(i,t)^2\}}$$

where $\Delta = \Delta_D \Delta_{L_h}$ combines the deconvolution and the differentiation operation, and $\Delta_{Lap}$ defines the 3D second-order difference operator. The spatial and temporal regularization parameters are given by the $\lambda_S$ and $\lambda_T$ , respectively. $N_i$ and $N_t$ denote the total number of voxels and timepoints, while $S(i)$ denotes all voxels in the neighborhood of voxel $i$. For more details about the implementation of the TA paradigm, we refer to (Karahanoğlu et al., 2013) and (Farouj et al., 2017), respectively.

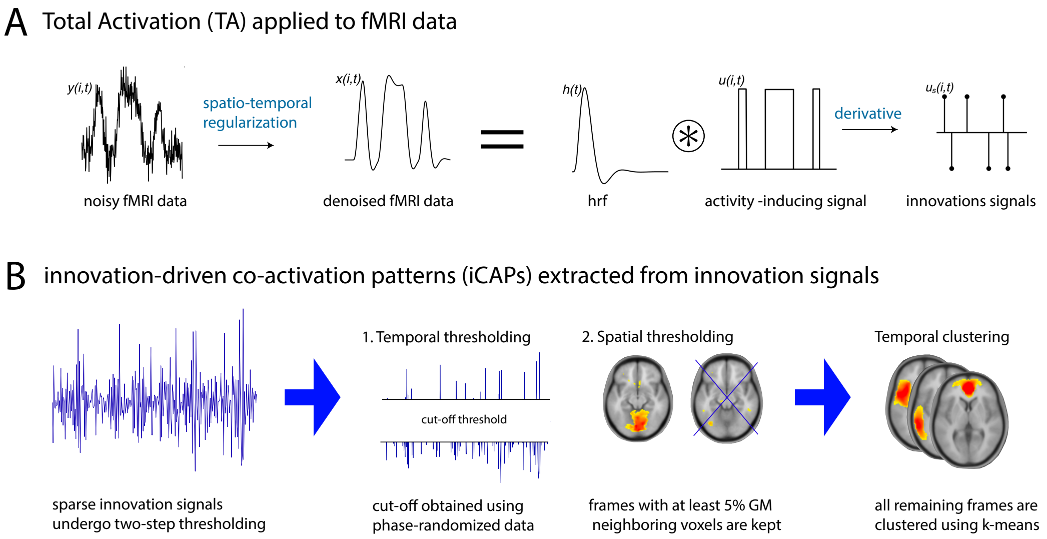


Fig. SI 1. **Methodological pipeline.** (A) Noisy fMRI time-courses are denoised using a combined spatio-temporal regularization, followed by a deconvolution from the HRF to obtain block-type activity inducing signals, which is then differentiated to get sparse innovation signals. (B) The resulting innovation signals undergo a two-step thresholding (spatial and temporal), and the remaining frames undergo temporal clustering to extract the iCAPs.

*Extraction of innovation-driven co-activation patterns (iCAPs)*

After running TA, we extract the sparse innovation signals, $u_S(i,t)$ or the *transients* by computing the temporal derivative of the activity-inducing signals $u(i,t)$. Then, the significant innovations, or *significant transients* are extracted by taking only a fraction across brain volumes following a two-step thresholding procedure described in Fig. SI 1. To determine the temporal threshold, the same dataset undergoes phase-randomization, from which we extract the cut-off threshold using the lowest 5th and highest 95th percentile of its corresponding innovation signal. Following the temporal thresholding, we take frames with at least 5% of the total number of gray matter voxels to undergo temporal clustering using k-means. The resulting maps, termed innovation-driven co-activation patterns (iCAPs) are shown in Fig. 2 in the main manuscript. For more details of the iCAPs extraction and the optimization of the thresholding, we refer to (Karahanoğlu and Van De Ville, 2015).

*Temporal Characteristics of iCAPs*

Finally, iCAP time-courses for each subject are computed using transient-informed spatio-temporal back-projection of the iCAP maps onto the activity-inducing signals (Zoller et al., 2019). The clustering of the innovation signals or the transient points for which we observe changes in neural activity allows for extraction of spatially and temporally overlapping spatial maps. This characteristic gives rise to a dynamically rich repertoire of functional states, each of which can be explored to observe iCAP-specific temporal characteristics.

*Finding the optimal number of clusters*

We used consensus clustering (Monti et al., 2003) to obtain the optimal number of clusters in the concatenated significant innovation frames. The method involved subsampling of the data and multiple runs of the clustering algorithm. The consistency of each frame to be grouped in a similar cluster is monitored through a consensus metric. Fig. SI 2 shows the consensus clustering matrices for all K values evaluated. The cumulative distribution of this metric is displayed in Fig. SI 3. We chose K = 17 to be the optimal number of clusters by looking at the trend of the area under the curve (AUC) corresponding to the consensus clustering matrices. This optimal number also coincides with the K that has the highest cluster consensus in Fig. SI 3(C).

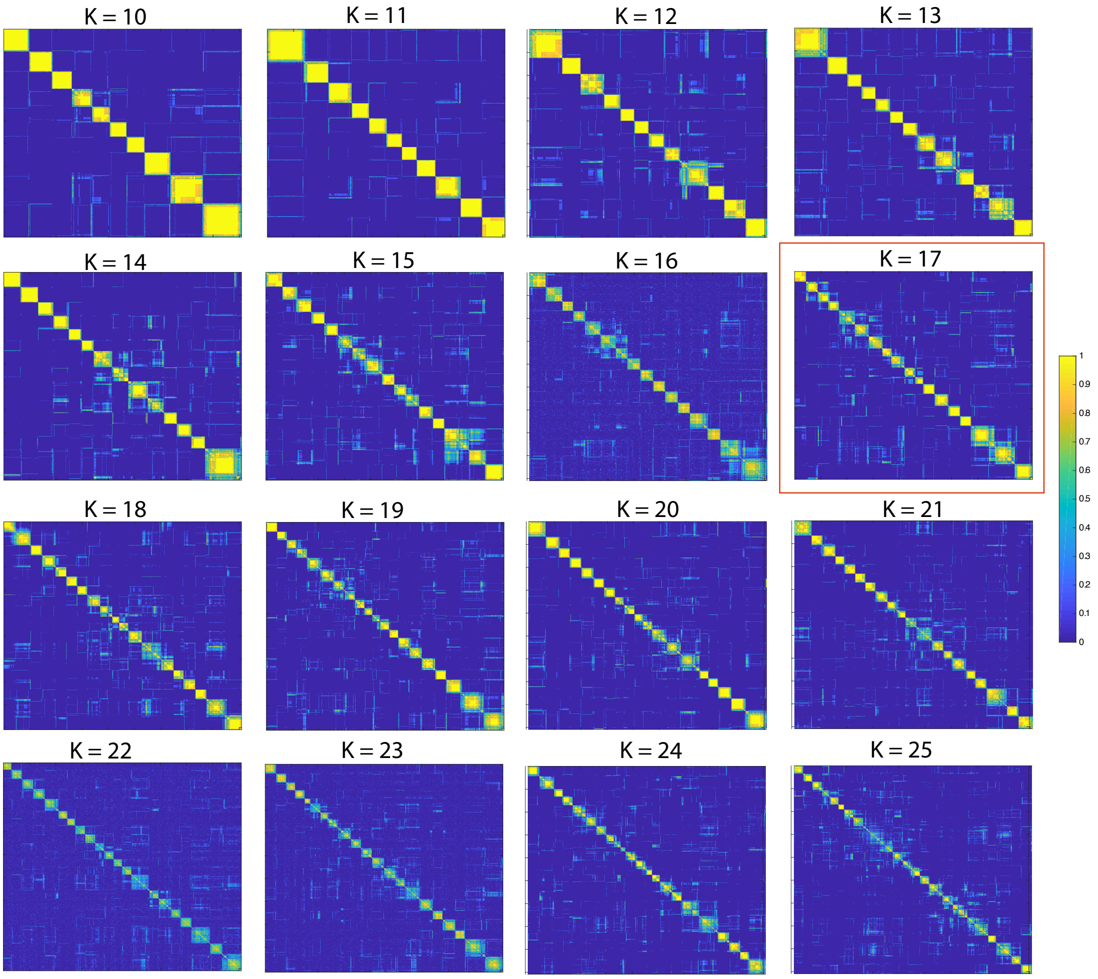


Fig. SI 2. **Consensus clustering matrices** for different cluster values K = {10, 11, 12, ..., 25}. The x and y axes correspond to frame number. Values in the matrix range from 0 to 1, which indicate the reproducibility of the sampling across multiple runs, with 1 being perfectly re-sampled at all times. Diagonal values are expected to be equal to 1 (the same frame indices will always be clustered into the same group). We chose K = 17 based on visual inspection and the consensus quality measures displayed in Fig. SI 3.

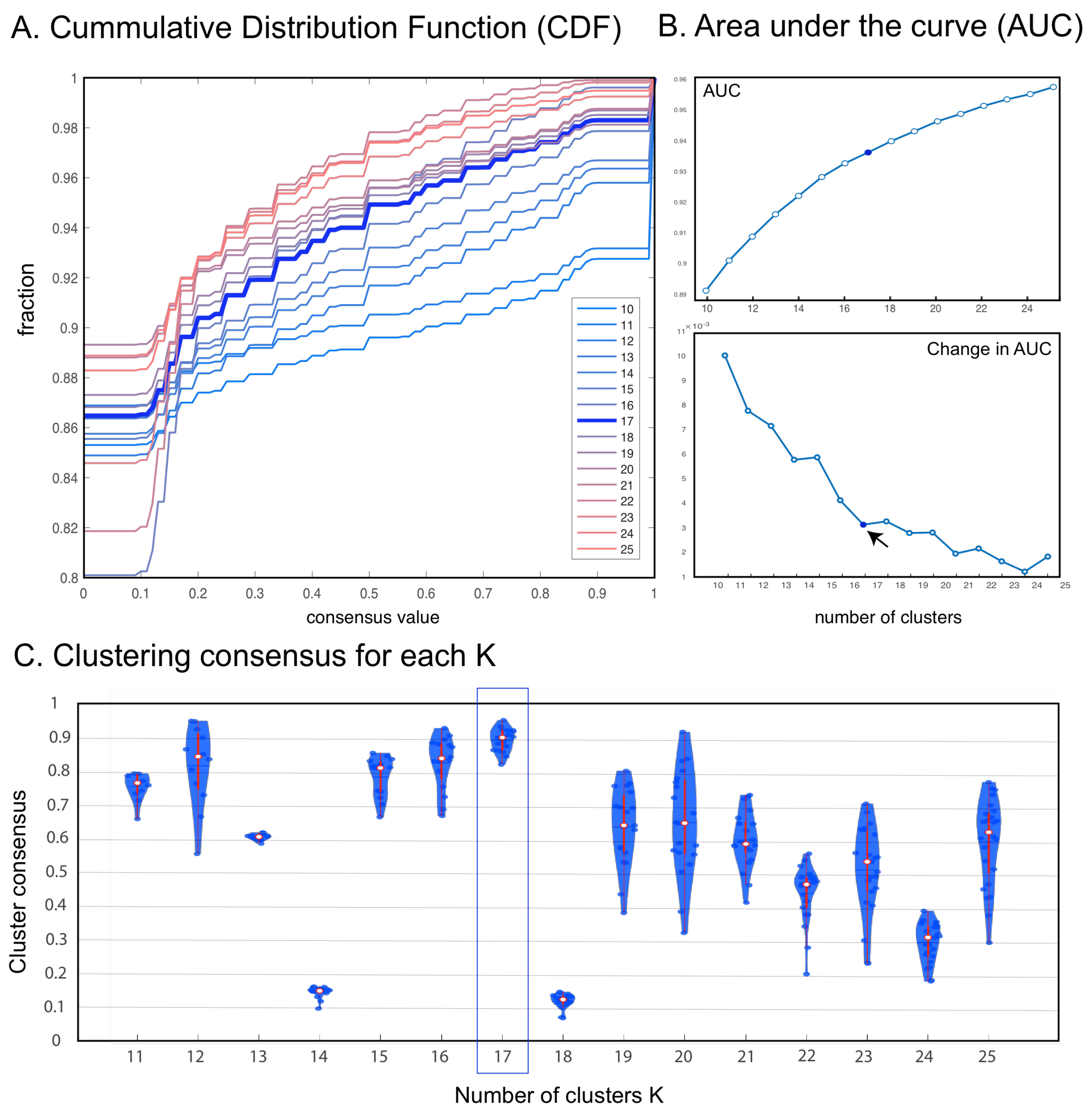


Fig. SI 3. **Consensus quality metrics.** The (A) cumulative distribution function (CDF) indicates the extent to which the consensus matrix distribution is skewed toward 0 and 1, with a flat line being the ideal shape (i.e., 0 means two frames are never clustered together while 1 means frames are always clustered together). The (B) area under the curve (AUC) of the CDF and the change in AUC displays the optimal number of cluster K to which there is minimal increase in the AUC. Finally, the (C) clustering consensus gives a measure of the stability of the observed iCAPs with respect to different K over multiple runs of the clustering; K=17 shows the highest cluster consensus. Using all these three consensus quality metrics, we chose K = 17 as the optimal number of clusters.

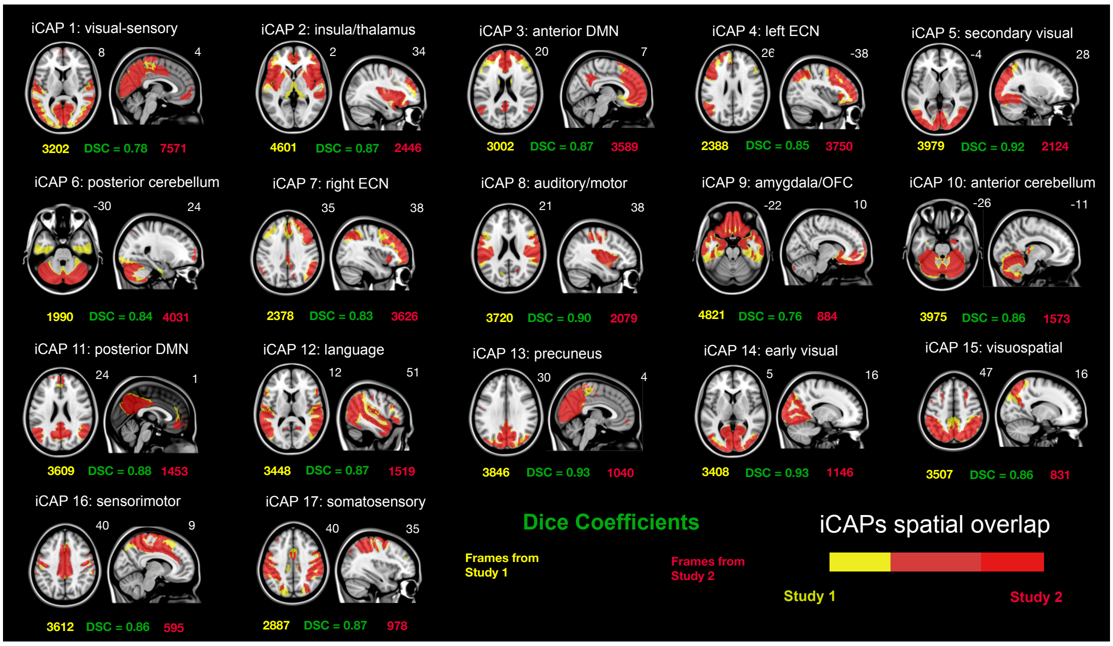


Fig. SI 4. **Spatial similarity** between iCAPs obtained by averaging the frame indices coming from Study 1 (yellow) versus iCAPs obtained by averaging the frame indices coming from Study 2 (red). The number of frames that contributed to the recovery of each iCAP coming from each dataset is written in red and yellow fonts. The iCAP maps are spatially z-scored and thresholded at $|z| > 1.5$. The Dice-Sorrensen Coefficient (DSC) is computed as a quantitative measure for the similarity of the two sets of iCAPs and is written in green font.

#### Supplementary Results

*Spatial characteristics of sleep-related iCAPs are robust across datasets*

In order to see whether the resulting iCAPs are robust with regards to the dataset from which the contributing frames belong to, we perform an averaging of significant frames corresponding to iCAP indices within a similar dataset (i.e., among those belonging Study 1 and among those belonging to Study 2). We computed the spatial similarity of the resulting iCAPs. Fig. SI 4 provides an overlay of the iCAPs computed using frames belonging to Study 1 versus Study 2. Visually, all iCAPs display similar spatial patterns in both the Study 1 and Study 2, which is also well supported by their Dice coefficients.

*Cluster assignments of significant innovations*

To obtain the iCAPs, we perform temporal clustering of significantly active frames. Significant frames from the Study 1 and Study 2 are concatenated together. This concatenated datamatrix is fed into the clustering algorithm, grouping together frames that are spatially similar. Study 1 contributes 58373 frames while the Study 2 contributes 39235, giving a total of 97608 significant frames. The number of contributing frames coming from each dataset is shown in Fig. SI 4 and is written in red and yellow fonts for Study 1 and Study 2, respectively. The distribution of clustering assignments is particularly telling, especially on the contribution of each dataset reflecting the tendency of the iCAP to appear in either Study 1 (reaching deep sleep) or in Study 2 (wake to N1). We observe the visual-sensory, anterior DMN, posterior cerebellum, and the left and right executive control networks to be mostly contributed by frames belonging to the Study 2 dataset. On the other hand, the Study 1 contributed more on the recovery of the remaining networks.

Table SI-1: iCAPs functional networks corresponding to the Greicius atlas (Shirer et al., 2012) and regions in the automated anatomical labeling atlas (Tzourio-Mazoyer et al., 2002). Percentiles indicate the fraction of voxels belonging to a network or region that has a z-score > 1.5. Only networks and regions that have at least 20% surviving percentile are included in the list. iCAP 6 is not included due to the excluded cerebellum in Greicius atlas. Specific regions that are not part of the cerebellum but are co-activated with iCAP 10 (anterior cerebellum) are included.

| ICAP | GREICIUS NETWORK ( %) | AAL | AAL REGION ( %) | Z-SCORE | VOXEL |
|---|---|---|---|---|---|
| 1 | Primary Visual (93.12 %) | Occipital | Cuneus L (83.29 %) | 2.35 | 289 |
| | Higher Visual (41.53 %) | Occipital | Calcarine L (72.69 %) | 2.41 | 386 |
| | Ventral DMN (41.06 %) | Occipital | Cuneus R (70.96 %) | 2.28 | 259 |
| | Precuneus (38.27 %) | Occipital | Calcarine R (70.33 %) | 2.48 | 294 |
| | | Occipital | Occipital Mid R (53.62 %) | 1.99 | 252 |
| | | Occipital | Lingual R (50.94 %) | 2.17 | 271 |
| | | Occipital | Occipital Inf L (49.77 %) | 2.07 | 110 |
| | | Occipital | Occipital Sup L (48.02 %) | 2.03 | 121 |
| | | Occipital | Occipital Mid L (47.02 %) | 2.1 | 363 |
| | | Occipital | Lingual L (46.35 %) | 2.15 | 254 |
| | | Occipital | Occipital Inf R (44.66 %) | 2.03 | 113 |
| | | Parietal | Precuneus R (42.88 %) | 2.04 | 304 |
| | | Parietal | Precuneus L (39.09 %) | 2.02 | 310 |
| | | Occipital | Occipital Sup R (38.25 %) | 1.92 | 109 |
| | | Parietal | Paracentral Lobule R (25.95 %) | 1.8 | 34 |
| | | Temporal | Temporal Sup L (21.65 %) | 1.78 | 118 |
| | | Occipital | Fusiform R (20.11 %) | 2 | 142 |
| 2 | Auditory (77.31 %) | Temporal | Heschl R (98.41 %) | 2.15 | 62 |
| | Basal Ganglia (52.46 %) | Limbic | Insula R (96.03 %) | 3.12 | 435 |
| | Anterior Salience (41.40 %) | Limbic | Insula L (93.74 %) | 2.89 | 464 |
| | Posterior Salience (23.94 %) | Temporal | Heschl L (89.86 %) | 1.98 | 62 |
| | | Central | Rolandic Oper L (86.75 %) | 2.23 | 216 |
| | | Central | Rolandic Oper R (86.52 %) | 2.23 | 276 |
| | | Subcortical | Putamen R (86.42 %) | 2.29 | 210 |
| | | Frontal | Frontal Inf Tri R (81.98 %) | 2.17 | 314 |
| | | Subcortical | Putamen L (77.69 %) | 2.18 | 188 |
| | | Frontal | Frontal Inf Oper R (77.27 %) | 2.51 | 238 |
| | | Frontal | Frontal Inf Oper L (75.69 %) | 2.19 | 165 |
| | | Subcortical | Pallidum R (75.00 %) | 2.01 | 9 |
| | | Frontal | Frontal Inf Tri L (49.48 %) | 2.04 | 239 |
| | | Temporal | Temporal Sup L (47.71 %) | 1.93 | 260 |
| | | Limbic | Cingulum Ant R (44.21 %) | 1.74 | 149 |
| | | Subcortical | Pallidum L (43.48 %) | 1.96 | 10 |
| | | Frontal | Frontal Inf Orb R (40.23 %) | 2.09 | 138 |
| | | Temporal | Temporal Pole Sup R (38.46 %) | 2.1 | 95 |
| | | Limbic | Cingulum Ant L (37.53 %) | 1.68 | 137 |
| | | Temporal | Temporal Sup R (32.11 %) | 1.94 | 228 |
| | | Subcortical | Thalamus R (27.50 %) | 1.63 | 44 |
| | | Temporal | Temporal Pole Sup L (27.27 %) | 2.03 | 63 |
| | | Frontal | Frontal Inf Orb L (23.97 %) | 2.09 | 93 |
| | | Parietal | SupraMarginal R (23.13 %) | 1.69 | 102 |
| | | Limbic | Cingulum Mid R (22.46 %) | 1.82 | 117 |
| | | Limbic | Amygdala R (21.88 %) | 1.69 | 14 |
| 3 | Dorsal DMN (76.38 %) | Frontal | Frontal Sup Orb Medial L | 3.51 | 160 |
| | Anterior Salience (35.23 %) | Frontal | Frontal Sup Medial L (91.98 %) | 3.27 | 436 |
| | | Frontal | Frontal Sup Orb Medial R | 3.46 | 197 |
| | | Frontal | Frontal Sup Medial R (91.08 | 3.14 | 398 |
| | | Limbic | Cingulum Ant R (90.21 %) | 3.57 | 304 |
| | | Limbic | Cingulum Ant L (89.86 %) | 3.61 | 328 |
| | | Limbic | Cingulum Post L (59.30 %) | 1.9 | 51 |
| | | Frontal | Frontal Sup L (57.89 %) | 2.3 | 356 |
| | | Frontal | Frontal Sup R (49.35 %) | 2.25 | 344 |

| | | | | | |
|---|---|---|---|---|---|
| | | Limbic | Cingulum Post R (39.39 %) | 1.94 | 13 |
| | | Frontal | Rectus L (35.62 %) | 2.37 | 78 |
| | | Frontal | Frontal Mid L (29.92 %) | 1.92 | 289 |
| | | Frontal | Rectus R (28.80 %) | 2.41 | 55 |
| | | Frontal | Frontal Sup Orb R (24.48 %) | 1.85 | 59 |
| | | Frontal | Frontal Sup Orb L (24.44 %) | 1.9 | 55 |
| | | Parietal | Angular L (23.59 %) | 1.7 | 67 |
| | | Limbic | Cingulum Mid R (20.73 %) | 2.05 | 108 |
| **4** | Left ECN (85.35 %) | Frontal | Frontal Inf Tri L (94.62 %) | 3.5 | 457 |
| | Language (51.24 %) | Parietal | Angular L (93.31 %) | 2.79 | 265 |
| | Visuospatial (31.20 %) | Frontal | Frontal Inf Oper L (89.91 %) | 3.32 | 196 |
| | Anterior Salience (24.77 %) | Frontal | Frontal Mid L (81.37 %) | 2.6 | 786 |
| | | Parietal | Parietal Inf L (66.50 %) | 2.69 | 397 |
| | | Frontal | Frontal Sup L (59.84 %) | 2.12 | 368 |
| | | Frontal | Frontal Inf Orb L (47.16 %) | 2.61 | 183 |
| | | Frontal | Precentral L (43.59 %) | 3.2 | 245 |
| | | Frontal | Frontal Sup Medial L (41.98 %) | 2.25 | 199 |
| | | Temporal | Temporal Mid L (39.00 %) | 2.05 | 454 |
| | | Frontal | Frontal Mid Orb L (33.85 %) | 2.67 | 66 |
| | | Frontal | Supp Motor Area L (33.03 %) | 2.4 | 143 |
| | | Parietal | Parietal Sup L (20.24 %) | 2.29 | 86 |
| **5** | Higher Visual (100.00 %) | Occipital | Occipital Inf R (100.00 %) | 3.56 | 253 |
| | | Occipital | Occipital Inf L (93.21 %) | 3.19 | 206 |
| | | Occipital | Occipital Sup R (84.56 %) | 2.83 | 241 |
| | | Occipital | Occipital Mid R (81.49 %) | 3.63 | 383 |
| | | Occipital | Occipital Mid L (77.46 %) | 3.49 | 598 |
| | | Occipital | Occipital Sup L (76.19 %) | 2.69 | 192 |
| | | Occipital | Lingual R (46.62 %) | 2.48 | 248 |
| | | Occipital | Calcarine L (46.14 %) | 2.5 | 245 |
| | | Occipital | Cuneus R (44.93 %) | 2.19 | 164 |
| | | Occipital | Lingual L (39.60 %) | 2.58 | 217 |
| | | Occipital | Cuneus L (38.90 %) | 2.12 | 135 |
| | | Occipital | Calcarine R (34.21 %) | 2.69 | 143 |
| | | Occipital | Fusiform R (30.31 %) | 2.4 | 214 |
| | | Occipital | Fusiform L (22.10 %) | 2.48 | 143 |
| **7** | Right ECN (67.49 %) | Frontal | Frontal Mid R (89.34 %) | 3.05 | 964 |
| | Precuneus (25.98 %) | Parietal | Angular R (81.94 %) | 2.96 | 295 |
| | Posterior Salience (23.67 %) | Parietal | Parietal Inf R (79.42 %) | 3.7 | 274 |
| | Left ECN (23.12 %) | Frontal | Frontal Mid Orb R (71.09 %) | 2.66 | 150 |
| | Anterior Salience (21.15 %) | Frontal | Frontal Inf Tri R (69.19 %) | 2.37 | 265 |
| | | Frontal | Frontal Sup R (64.56 %) | 2.41 | 450 |
| | | Frontal | Frontal Inf Oper R (62.01 %) | 2.28 | 191 |
| | | Frontal | Frontal Sup Medial R (43.02 | 2.19 | 188 |
| | | Parietal | SupraMarginal R (36.05 %) | 2.76 | 159 |
| | | Frontal | Frontal Inf Orb R (34.69 %) | 2.38 | 119 |
| | | Frontal | Frontal Sup Orb R (29.05 %) | 2.55 | 70 |
| | | Parietal | Parietal Inf L (28.48 %) | 1.99 | 170 |
| | | Limbic | Cingulum Ant R (27.30 %) | 1.91 | 92 |
| | | Limbic | Cingulum Mid R (25.53 %) | 2 | 133 |
| | | Frontal | Frontal Mid Orb L (23.59 %) | 1.95 | 46 |
| | | Parietal | Angular L (23.24 %) | 1.84 | 66 |
| **8** | Auditory (94.91 %) | Temporal | Heschl L (100.00 %) | 3.13 | 69 |
| | Posterior Salience (45.88 %) | Temporal | Heschl R (100.00 %) | 3.48 | 63 |
| | Sensorimotor (28.50 %) | Central | Rolandic Oper R (96.87 %) | 3.35 | 309 |
| | | Central | Rolandic Oper L (94.78 %) | 2.8 | 236 |
| | | Parietal | SupraMarginal L (74.56 %) | 2.6 | 214 |
| | | Temporal | Temporal Sup L (72.48 %) | 2.69 | 395 |
| | | Parietal | Postcentral L (59.46 %) | 3.38 | 396 |
| | | Temporal | Temporal Sup R (57.61 %) | 2.74 | 409 |
| | | Parietal | SupraMarginal R (57.60 %) | 2.97 | 254 |
| | | Parietal | Postcentral R (51.89 %) | 3.38 | 343 |
| | | Frontal | Precentral R (47.22 %) | 2.58 | 246 |
| | | Limbic | Insula R (46.36 %) | 2.39 | 210 |
| | | Limbic | Insula L (33.33 %) | 2.29 | 165 |
| | | Frontal | Precentral L (22.60 %) | 2.44 | 127 |
| **9** | | Frontal | Rectus L (100.00 %) | 3.13 | 219 |
| | | Frontal | Rectus R (100.00 %) | 3.05 | 191 |
| | | Frontal | Olfactory R (97.30 %) | 2.7 | 72 |
| | | Frontal | Olfactory L (96.00 %) | 2.81 | 72 |
| | | Temporal | Temporal Pole Mid L (94.84 | 2.33 | 147 |

| | | | | | |
|---|---|---|---|---|---|
| | | Frontal | Frontal Sup Orb L (94.22 %) | 2.47 | 212 |
| | | Frontal | Frontal Sup Orb R (94.19 %) | 2.5 | 227 |
| | | Temporal | Temporal Pole Mid R (90.87 | 2.04 | 189 |
| | | Limbic | Amygdala L (89.66 %) | 2.07 | 52 |
| | | Frontal | Frontal Sup Orb Medial L | 2.12 | 145 |
| | | Frontal | Frontal Mid Orb R (86.73 %) | 2.54 | 183 |
| | | Frontal | Frontal Inf Orb L (86.34 %) | 2.44 | 335 |
| | | Limbic | Amygdala R (85.94 %) | 1.94 | 55 |
| | | Frontal | Frontal Mid Orb L (84.62 %) | 2.49 | 165 |
| | | Temporal | Temporal Pole Sup L (82.68 %) | 2.17 | 191 |
| | | Frontal | Frontal Sup Orb Medial R | 2.13 | 176 |
| | | Frontal | Frontal Inf Orb R (76.68 %) | 2.3 | 263 |
| | | Temporal | Temporal Pole Sup R (75.71 %) | 2.03 | 187 |
| | | Limbic | ParaHippocampal R (69.61 %) | 1.98 | 197 |
| | | Limbic | ParaHippocampal L (69.39 %) | 2.03 | 170 |
| | | Temporal | Temporal Inf L (55.67 %) | 1.86 | 447 |
| | | Limbic | Hippocampus L (53.60 %) | 2.03 | 119 |
| | | Limbic | Hippocampus R (48.23 %) | 1.88 | 109 |
| | | Temporal | Temporal Inf R (46.24 %) | 1.81 | 406 |
| | | Subcortical | Caudate L (37.43 %) | 2.17 | 70 |
| | | Subcortical | Caudate R (36.04 %) | 2.13 | 71 |
| | | Occipital | Fusiform L (31.22 %) | 1.84 | 202 |
| | | Occipital | Fusiform R (29.18 %) | 1.75 | 206 |
| | | Temporal | Temporal Mid L (25.26 %) | 1.85 | 294 |
| **10** | Sensorimotor (50.65 %) | Occipital | Fusiform R (59.07 %) | 2.55 | 417 |
| | | Occipital | Fusiform L (54.71 %) | 2.34 | 354 |
| | | Occipital | Lingual R (41.92 %) | 2.22 | 223 |
| | | Occipital | Lingual L (39.42 %) | 2.19 | 216 |
| | | Limbic | ParaHippocampal R (27.56 %) | 2.13 | 78 |
| | | Limbic | ParaHippocampal L (24.90 %) | 1.91 | 61 |
| **11** | Precuneus (84.88 %) | Parietal | Angular L (97.89 %) | 2.86 | 278 |
| | Ventral DMN (50.52 %) | Limbic | Cingulum Post L (97.67 %) | 4.71 | 84 |
| | Left ECN (30.98 %) | Limbic | Cingulum Post R (93.94 %) | 4.28 | 31 |
| | Dorsal DMN (26.26 %) | Parietal | Angular R (88.61 %) | 2.82 | 319 |
| | Language (23.04 %) | Parietal | Precuneus R (72.78 %) | 3.84 | 516 |
| | | Parietal | Precuneus L (64.82 %) | 4.13 | 514 |
| | | Parietal | Parietal Inf R (33.62 %) | 2.32 | 116 |
| | | Occipital | Cuneus L (32.85 %) | 2.83 | 114 |
| | | Limbic | Cingulum Mid L (30.71 %) | 3.1 | 156 |
| | | Limbic | Cingulum Mid R (24.95 %) | 3.51 | 130 |
| | | Occipital | Cuneus R (24.38 %) | 2.29 | 89 |
| | | Parietal | Parietal Inf L (21.44 %) | 2.18 | 128 |
| **12** | Language (74.17 %) | Parietal | Angular L (86.97 %) | 2.68 | 247 |
| | Auditory (51.16 %) | Temporal | Temporal Sup R (82.68 %) | 3.14 | 587 |
| | Posterior Salience (24.87 %) | Temporal | Temporal Mid R (82.56 %) | 3.11 | 876 |
| | | Temporal | Temporal Mid L (77.23 %) | 3.05 | 899 |
| | | Temporal | Temporal Sup L (66.06 %) | 2.35 | 360 |
| | | Parietal | Angular R (62.22 %) | 2.8 | 224 |
| | | Parietal | SupraMarginal L (37.98 %) | 2.44 | 109 |
| | | Parietal | SupraMarginal R (31.29 %) | 2.51 | 138 |
| | | Parietal | Parietal Inf R (21.45 %) | 2.11 | 74 |
| **13** | Primary Visual (100.00 %) | Occipital | Cuneus L (97.69 %) | 3.82 | 339 |
| | Precuneus (60.79 %) | Occipital | Cuneus R (89.32 %) | 3.63 | 326 |
| | Ventral DMN (24.59 %) | Occipital | Calcarine R (81.10 %) | 4.06 | 339 |
| | | Occipital | Calcarine L (68.55 %) | 4.05 | 364 |
| | | Occipital | Lingual R (56.95 %) | 3.06 | 303 |
| | | Occipital | Lingual L (49.82 %) | 2.88 | 273 |
| | | Parietal | Precuneus R (48.94 %) | 2.88 | 347 |
| | | Parietal | Precuneus L (36.19 %) | 2.68 | 287 |
| | | Occipital | Occipital Sup L (34.13 %) | 2.31 | 86 |
| | | Limbic | Cingulum Post L (31.40 %) | 1.74 | 27 |
| | | Occipital | Occipital Sup R (22.46 %) | 1.93 | 64 |
| | | Limbic | Cingulum Post R (21.21 %) | 1.64 | 7 |
| **14** | Primary Visual (100.00 %) | Occipital | Calcarine L (90.96 %) | 3.83 | 483 |
| | Higher Visual (48.91 %) | Occipital | Calcarine R (88.52 %) | 3.98 | 370 |
| | | Occipital | Cuneus R (85.75 %) | 3.28 | 313 |
| | | Occipital | Cuneus L (83.57 %) | 3.58 | 290 |
| | | Occipital | Lingual R (81.39 %) | 2.96 | 433 |
| | | Occipital | Occipital Sup L (71.83 %) | 2.73 | 181 |
| | | Occipital | Lingual L (67.15 %) | 3.04 | 368 |

| | | | | | |
|---|---|---|---|---|---|
| | | Occipital | Occipital Sup R (61.75 %) | 2.88 | 176 |
| | | Occipital | Occipital Mid R (37.66 %) | 1.91 | 177 |
| | | Occipital | Occipital Mid L (29.27 %) | 1.95 | 226 |
| **15** | Precuneus (59.84 %) | Parietal | Parietal Sup L (87.76 %) | 3.88 | 373 |
| | Visuospatial (47.29 %) | Parietel | Parietal Sup R (86.87 %) | 3.58 | 311 |
| | Ventral DMN (43.44 %) | Parietal | Parietal Inf R (82.61 %) | 2.85 | 285 |
| | Left ECN (27.89 %) | Parietal | Parietal Inf L (71.86 %) | 3 | 429 |
| | | Parietal | Angular R (53.61 %) | 3.21 | 193 |
| | | Occipital | Occipital Sup R (48.07 %) | 3.67 | 137 |
| | | Occipital | Occipital Sup L (47.22 %) | 2.8 | 119 |
| | | Parietal | Precuneus L (46.53 %) | 2.75 | 369 |
| | | Occipital | Occipital Mid R (44.68 %) | 2.6 | 210 |
| | | Parietal | Precuneus R (41.75 %) | 2.61 | 296 |
| | | Occipital | Occipital Mid L (32.38 %) | 2.56 | 250 |
| | | Parietal | Angular L (30.99 %) | 2.1 | 88 |
| **16** | Sensorimotor (34.46 %) | Parietal | Paracentral Lobule L (100.00 | 4.42 | 166 |
| | Anterior Salience (32.92 %) | Parietal | Paracentral Lobule R (100.00 | 4.37 | 131 |
| | | Frontal | Supp Motor Area R (86.04 %) | 3.59 | 413 |
| | | Limbic | Cingulum Mid L (78.74 %) | 2.87 | 400 |
| | | Frontal | Supp Motor Area L (75.98 %) | 3.25 | 329 |
| | | Limbic | Cingulum Mid R (67.37 %) | 2.99 | 351 |
| | | Frontal | Precentral R (36.08 %) | 2.37 | 188 |
| | | Parietal | Postcentral R (33.13 %) | 2.73 | 219 |
| | | Parietal | Postcentral L (29.58 %) | 2.32 | 197 |
| | | Parietal | Precuneus L (27.99 %) | 3.02 | 222 |
| | | Frontal | Precentral L (25.98 %) | 2.39 | 146 |
| | | Frontal | Frontal Sup R (25.68 %) | 2.33 | 179 |
| | | Parietal | Precuneus R (23.55 %) | 2.61 | 167 |
| | | Parietel | Parietal Sup R (20.11 %) | 2.21 | 72 |
| **17** | Visuospatial (58.76 %) | Parietel | Parietal Sup R (76.26 %) | 3.01 | 273 |
| | Posterior Salience (51.33 %) | Parietal | Postcentral R (68.68 %) | 3.83 | 454 |
| | Sensorimotor (34.46 %) | Parietal | SupraMarginal R (64.85 %) | 2.79 | 286 |
| | | Parietal | Parietal Sup L (64.00 %) | 2.96 | 272 |
| | | Parietal | Parietal Inf L (60.80 %) | 3.39 | 363 |
| | | Parietal | Postcentral L (59.46 %) | 3.28 | 396 |
| | | Parietal | Parietal Inf R (55.36 %) | 3.43 | 191 |
| | | Parietal | SupraMarginal L (47.39 %) | 2.75 | 136 |
| | | Frontal | Precentral R (45.87 %) | 2.26 | 239 |
| | | Frontal | Precentral L (40.21 %) | 2.23 | 226 |
| | | Frontal | Frontal Sup R (20.66 %) | 2.28 | 144 |

Table SI-2: Test statistics corresponding to Figure 3 (A) and (C) of the main manuscript. Results from two-sample paired t-tests and 1000 rounds of permutation testing. We compared iCAPs RCD and average durations across different sleep stages. Effect size corresponds to Cohen's d statistic.

| iCAP | | Wake-N1 | Wake-N2 | Wake-N3 | N1-N2 | N1-N3 | N2-N3 |
|---|---|---|---|---|---|---|---|
| **1. visual-sensory** | | | | | | | |
| RCD | t-statistic | -3.92 | -6.22 | 1.69 | -2.3 | 5.61 | 7.91 |
| | p-value | 0.357 | 0.154 | 0.715 | 0.511 | 0.122 | 0.025 |
| | Effect-size | -0.34 | -0.57 | 0.15 | -0.25 | 0.6 | 0.88 |
| average durations | t-statistic | -3.92 | -6.22 | 1.69 | -2.3 | 5.61 | 7.91 |
| | p-value | 0.357 | 0.154 | 0.715 | 0.511 | 0.122 | 0.025 |
| | Effect-size | -0.34 | -0.57 | 0.15 | -0.25 | 0.6 | 0.88 |
| **2. insula/thalamus** | | | | | | | |
| RCD | t-statistic | -0.01 | -7.8 | 7.99 | -7.79 | 8 | 15.79 |
| | p-value | 0.999 | 0.011 | 0.031 | 0.031 | 0.042 | 0.002 |
| | Effect-size | 0 | -1.01 | 0.92 | -0.87 | 0.8 | 1.71 |
| average durations | t-statistic | -0.01 | -7.8 | 7.99 | -7.79 | 8 | 15.79 |
| | p-value | 0.999 | 0.011 | 0.031 | 0.031 | 0.042 | 0.002 |
| | Effect-size | 0 | -1.01 | 0.92 | -0.87 | 0.8 | 1.71 |
| **3. anterior DMN** | | | | | | | |
| RCD | t-statistic | 6.97 | 1.79 | 15.63 | -5.17 | 8.66 | 13.83 |
| | p-value | 0.07 | 0.581 | 0.001 | 0.063 | 0.009 | 0.001 |
| | Effect-size | 0.74 | 0.23 | 1.81 | -0.73 | 1.1 | 2.21 |
| average durations | t-statistic | 6.97 | 1.79 | 15.63 | -5.17 | 8.66 | 13.83 |
| | p-value | 0.07 | 0.581 | 0.001 | 0.063 | 0.009 | 0.001 |
| | Effect-size | 0.74 | 0.23 | 1.81 | -0.73 | 1.1 | 2.21 |
| **4. left ECN** | | | | | | | |

| | | | | | | | |
|---|---|---|---|---|---|---|---|
| RCD | t-statistic | 4.4 | 3.68 | 15.34 | -0.73 | 10.94 | 11.66 |
| | p-value | 0.16 | 0.119 | 0.001 | 0.761 | 0.002 | 0.001 |
| | Effect-size | 0.51 | 0.66 | 1.91 | -0.12 | 1.31 | 2.25 |
| average durations | t-statistic | 4.4 | 3.68 | 15.34 | -0.73 | 10.94 | 11.66 |
| | p-value | 0.16 | 0.119 | 0.001 | 0.761 | 0.002 | 0.001 |
| | Effect-size | 0.51 | 0.66 | 1.91 | -0.12 | 1.31 | 2.25 |
| **5. secondary visual** | | | | | | | |
| RCD | t-statistic | -5.79 | -6.72 | -2.34 | -0.93 | 3.44 | 4.38 |
| | p-value | 0.103 | 0.005 | 0.364 | 0.797 | 0.354 | 0.139 |
| | Effect-size | -0.74 | -1.23 | -0.36 | -0.11 | 0.35 | 0.59 |
| average durations | t-statistic | -5.79 | -6.72 | -2.34 | -0.93 | 3.44 | 4.38 |
| | p-value | 0.103 | 0.005 | 0.364 | 0.797 | 0.354 | 0.139 |
| | Effect-size | -0.74 | -1.23 | -0.36 | -0.11 | 0.35 | 0.59 |
| **6. posterior CEB** | | | | | | | |
| RCD | t-statistic | -0.21 | 2.76 | 7.88 | 2.97 | 8.09 | 5.12 |
| | p-value | 0.951 | 0.356 | 0.002 | 0.344 | 0.008 | 0.016 |
| | Effect-size | -0.02 | 0.4 | 1.15 | 0.41 | 1.12 | 1.02 |
| average durations | t-statistic | -0.21 | 2.76 | 7.88 | 2.97 | 8.09 | 5.12 |
| | p-value | 0.951 | 0.356 | 0.002 | 0.344 | 0.008 | 0.016 |
| | Effect-size | -0.02 | 0.4 | 1.15 | 0.41 | 1.12 | 1.02 |
| **7. right ECN** | | | | | | | |
| RCD | t-statistic | 5.45 | 3.64 | 18.2 | -1.81 | 12.75 | 14.56 |
| | p-value | 0.082 | 0.22 | 0.001 | 0.316 | 0.001 | 0.001 |
| | Effect-size | 0.71 | 0.55 | 2.03 | -0.39 | 1.84 | 2.47 |
| average durations | t-statistic | 5.45 | 3.64 | 18.2 | -1.81 | 12.75 | 14.56 |
| | p-value | 0.082 | 0.22 | 0.001 | 0.316 | 0.001 | 0.001 |
| | Effect-size | 0.71 | 0.55 | 2.03 | -0.39 | 1.84 | 2.47 |
| **8. auditory/motor** | | | | | | | |
| RCD | t-statistic | -2.07 | -7.61 | -2.68 | -5.54 | -0.61 | 4.93 |
| | p-value | 0.308 | 0.003 | 0.22 | 0.034 | 0.804 | 0.069 |
| | Effect-size | -0.39 | -1.31 | -0.48 | -0.86 | -0.1 | 0.73 |
| average durations | t-statistic | -2.07 | -7.61 | -2.68 | -5.54 | -0.61 | 4.93 |
| | p-value | 0.308 | 0.003 | 0.22 | 0.034 | 0.804 | 0.069 |
| | Effect-size | -0.39 | -1.31 | -0.48 | -0.86 | -0.1 | 0.73 |
| **9. amygdala/OFC** | | | | | | | |
| RCD | t-statistic | 0.48 | -2.06 | 7.45 | -2.54 | 6.97 | 9.51 |
| | p-value | 0.885 | 0.552 | 0.022 | 0.481 | 0.03 | 0.001 |
| | Effect-size | 0.05 | -0.23 | 0.89 | -0.29 | 0.86 | 1.25 |
| average durations | t-statistic | 0.48 | -2.06 | 7.45 | -2.54 | 6.97 | 9.51 |
| | p-value | 0.885 | 0.552 | 0.022 | 0.481 | 0.03 | 0.001 |
| | Effect-size | 0.05 | -0.23 | 0.89 | -0.29 | 0.86 | 1.25 |
| **10. anterior CEB** | | | | | | | |
| RCD | t-statistic | 1.09 | -2.28 | 7.46 | -3.38 | 6.36 | 9.74 |
| | p-value | 0.784 | 0.501 | 0.024 | 0.38 | 0.089 | 0.004 |
| | Effect-size | 0.11 | -0.26 | 0.93 | -0.34 | 0.7 | 1.18 |
| average durations | t-statistic | 1.09 | -2.28 | 7.46 | -3.38 | 6.36 | 9.74 |
| | p-value | 0.784 | 0.501 | 0.024 | 0.38 | 0.089 | 0.004 |
| | Effect-size | 0.11 | -0.26 | 0.93 | -0.34 | 0.7 | 1.18 |
| **11. posterior DMN** | | | | | | | |
| RCD | t-statistic | 0.62 | -5.07 | -2.82 | -5.69 | -3.44 | 2.25 |
| | p-value | 0.769 | 0.012 | 0.151 | 0.006 | 0.108 | 0.284 |
| | Effect-size | 0.12 | -1.01 | -0.56 | -1.06 | -0.63 | 0.43 |
| average durations | t-statistic | 0.62 | -5.07 | -2.82 | -5.69 | -3.44 | 2.25 |
| | p-value | 0.769 | 0.012 | 0.151 | 0.006 | 0.108 | 0.284 |
| | Effect-size | 0.12 | -1.01 | -0.56 | -1.06 | -0.63 | 0.43 |
| **12. language** | | | | | | | |
| RCD | t-statistic | -1.89 | -11.53 | -4.09 | -9.65 | -2.2 | 7.44 |
| | p-value | 0.532 | 0.002 | 0.217 | 0.009 | 0.581 | 0.05 |
| | Effect-size | -0.25 | -1.49 | -0.49 | -1.09 | -0.24 | 0.79 |
| average durations | t-statistic | -1.89 | -11.53 | -4.09 | -9.65 | -2.2 | 7.44 |
| | p-value | 0.532 | 0.002 | 0.217 | 0.009 | 0.581 | 0.05 |
| | Effect-size | -0.25 | -1.49 | -0.49 | -1.09 | -0.24 | 0.79 |
| **13. precuneus** | | | | | | | |
| RCD | t-statistic | -4.84 | -7.15 | -4.1 | -2.31 | 0.74 | 3.05 |
| | p-value | 0.13 | 0.036 | 0.14 | 0.516 | 0.79 | 0.356 |
| | Effect-size | -0.58 | -0.83 | -0.59 | -0.25 | 0.1 | 0.38 |
| average durations | t-statistic | -4.84 | -7.15 | -4.1 | -2.31 | 0.74 | 3.05 |
| | p-value | 0.13 | 0.036 | 0.14 | 0.516 | 0.79 | 0.356 |
| | Effect-size | -0.58 | -0.83 | -0.59 | -0.25 | 0.1 | 0.38 |
| **14. primary visual** | | | | | | | |
| RCD | t-statistic | -5.75 | -8.57 | -6.23 | -2.82 | -0.48 | 2.34 |
| | p-value | 0.025 | 0.002 | 0.008 | 0.313 | 0.839 | 0.343 |
| | Effect-size | -0.89 | -1.69 | -1.11 | -0.41 | -0.06 | 0.39 |
| average durations | t-statistic | -5.75 | -8.57 | -6.23 | -2.82 | -0.48 | 2.34 |
| | p-value | 0.025 | 0.002 | 0.008 | 0.313 | 0.839 | 0.343 |
| | Effect-size | -0.89 | -1.69 | -1.11 | -0.41 | -0.06 | 0.39 |
| **15. visuospatial** | | | | | | | |
| RCD | t-statistic | 0.75 | -4.59 | -0.04 | -5.34 | -0.79 | 4.55 |
| | p-value | 0.834 | 0.059 | 0.986 | 0.049 | 0.833 | 0.032 |
| | Effect-size | 0.09 | -0.79 | -0.01 | -0.8 | -0.11 | 0.87 |
| average durations | t-statistic | 0.75 | -4.59 | -0.04 | -5.34 | -0.79 | 4.55 |
| | p-value | 0.834 | 0.059 | 0.986 | 0.049 | 0.833 | 0.032 |

| | | | | | | | |
|---|---|---|---|---|---|---|---|
| | Effect-size | 0.09 | -0.79 | -0.01 | -0.8 | -0.11 | 0.87 |
| **16. sensorimotor** | | | | | | | |
| RCD | t-statistic | -0.74 | -14.78 | -9.18 | -14.04 | -8.44 | 5.6 |
| | p-value | 0.743 | 0.001 | 0.001 | 0.002 | 0.002 | 0.163 |
| | Effect-size | -0.14 | -1.71 | -1.43 | -1.62 | -1.32 | 0.57 |
| average durations | t-statistic | -0.74 | -14.78 | -9.18 | -14.04 | -8.44 | 5.6 |
| | p-value | 0.743 | 0.001 | 0.001 | 0.002 | 0.002 | 0.163 |
| | Effect-size | -0.14 | -1.71 | -1.43 | -1.62 | -1.32 | 0.57 |
| **17. somatosensory** | | | | | | | |
| RCD | t-statistic | -0.37 | -7.53 | -3.03 | -7.15 | -2.66 | 4.49 |
| | p-value | 0.854 | 0.004 | 0.228 | 0.014 | 0.355 | 0.107 |
| | Effect-size | -0.06 | -1.24 | -0.47 | -1.03 | -0.36 | 0.62 |
| average durations | t-statistic | -0.37 | -7.53 | -3.03 | -7.15 | -2.66 | 4.49 |
| | p-value | 0.854 | 0.004 | 0.228 | 0.014 | 0.355 | 0.107 |
| | Effect-size | -0.06 | -1.24 | -0.47 | -1.03 | -0.36 | 0.62 |

Table SI-3: Test statistics corresponding to Figure 4 (A) of the main manuscript. Results from ANOVA and the corrected p-values after multiple comparison test. Asterisks mark p-values less than 0.05.

| N | F-STATISTIC | PROB > F | WAKE - | WAKE - | WAKE- | N1 - N2 | N1 - N3 | N2 - N3 |
|---|---|---|---|---|---|---|---|---|
| **0** | 2.36 | 0.08 | 0.18 | 0.97 | 0.79 | 0.09 | 0.81 | 0.58 |
| **1** | 3.24 | 0.03 | 0.69 | 0.02* | 0.98 | 0.29 | 0.93 | 0.14 |
| **2** | 4.65 | 0.01 | 0.98 | 0.01* | 0.99 | 0.05* | 0.93 | 0.02* |
| **3** | 4.74 | 0.01 | 1 | 0.04* | 0.81 | 0.03* | 0.91 | 0.01* |
| **4** | 4.33 | 0.01 | 0.59 | 0.23 | 0.49 | 0.02* | 0.99 | 0.02* |
| **5** | 6.17 | 0 | 0.4 | 0.18 | 0.19 | 0.01* | 0.93 | 0* |
| **6** | 1.08 | 0.37 | 1 | 0.81 | 0.72 | 0.75 | 0.81 | 0.3 |
| **7** | 1.26 | 0.3 | 0.83 | 1 | 0.34 | 0.83 | 0.8 | 0.34 |
| **8** | 0.9 | 0.45 | 0.63 | 1 | 0.61 | 0.71 | 1 | 0.68 |
| **9** | 1.07 | 0.37 | 0.62 | 1 | 0.51 | 0.68 | 0.99 | 0.56 |
| **10** | 1.53 | 0.217 | 0.55 | 1 | 0.32 | 0.58 | 0.98 | 0.35 |

Table SI-4: Test statistics corresponding to Figure 4 (B) and (C) of the main manuscript. Results from ANOVA and the corrected p-values after multiple comparison test.

| | F- | PROB > | WAKE - | WAKE - | WAKE- | N1 - | N1 - | N2 - |
|---|---|---|---|---|---|---|---|---|
| **SAME-SIGNED** | 22.71 | 3.9E-10 | 4.5E-7 | 0.0091 | 3E-4 | 1.3E-9 | 0.009 | 4.6E-7 |
| **OPPOSITE-** | 12.92 | 1E-6 | 2.3E-6 | 0.0051 | 0.0012 | 7.1E-8 | 8.8E-4 | 1.2E-4 |
| **PEARSON** | 33.13 | 4.7E-13 | 0.2571 | 1.8E-7 | 3.3E-9 | 1.46-7 | 6.3E-9 | 0.115 |
